# Focused ultrasound neuromodulation of mediodorsal thalamus disrupts decision flexibility during reward learning

**DOI:** 10.1101/2025.06.03.657634

**Authors:** Graeme Mackenzie, William Gilmour, Isla Barnard, Shen-Shen Yang, Szabolcs Suveges, Jennifer MacFarlane, Avinash Kanodia, James Manfield, Sadaquate Khan, Jibril Osman-Farah, Antonella Macerollo, Douglas Steele, Tom Gilbertson

## Abstract

When learning to find the most beneficial course of action, the prefrontal cortex guides decisions by comparing estimates of the relative value of the options available. Basic neuroscience studies in animals support the view that the thalamus can regulate this activity within and across the prefrontal cortex. We studied a group of patients (n=37) undergoing unilateral MR guided focused ultrasound for essential tremor, performing the restless bandit, a reward reinforcement learning task, immediately before and after thalamotomy.

Thalamotomy significantly impaired the proportion of switch choices during the task without affecting overall performance. This effect was observed when the task was delivered to co-incide with maximal vasogenic thalamic oedema but not in a control group tested on the same day of their treatment. A reinforcement learning model fitted to the patients’ choices replicated the effect of thalamotomy when the model increased exploitation of the bandits’ learnt value estimate. This shift in the explore-exploit trade-off, manifesting as reduced choice flexibility, co-varied with the pattern of post-operative oedema extension into mediodorsal nucleus.

These findings confirm a causal role of the thalamus and specifically the mediodorsal nucleus, in regulating the extent to which value estimates are used to guide decisions and learning from reward.

## Introduction

Survival relies upon adaptation to environmental changes and making decisions that lead to maximum utility. In humans, the prefrontal cortex (PFC), consists of multiple sub-regions which contribute discrete functions to both behavioural control and motivation ^1–4^. These furnish the human brain with cognitive flexibility by allocating values to actions which are based upon learnt experience^3,5–7^. A feature of human decision making is the ability to dynamically alter the extent to which value estimates are incorporated and expressed into behaviour^8–10^. Neural activity in the PFC encodes behavioural strategies under conditions where environmental stability requires maintenance of the *status quo* to exploit confidence in the reliability of a learnt choices value^8,9,11–13^. Equally, when the correct course of action is unclear, activity in the PFC predicts adaptive, exploratory strategies^8,9,13^. Exploration aims to minimise uncertainty, and regain confidence in the reliability of the value-estimate, by updating the estimate through knowledge gained from experiencing an alternative course of action^11,14–17^. Evidence from the past two decades has converged on the existence of two partly distinct PFC circuits which contribute to these aspects of executive function^18^. Cognitive control, which imparts the ability to monitor and switch behaviour, inhibit actions, and detect conflict, has been associated with the dorsolateral PFC (DLPFC) and the anterior cingulate cortex (ACC)^19,20^. In contrast, a valuation network, centred on the ventromedial and lateral orbitofrontal regions (vmPFC/ OFC), guides decisions through learning and encoding neural estimates of decision value^21,22^. It is unclear how these circuits interact, and how their respective neural functions are integrated to promote switching between these types of behaviour. Converging evidence suggests subcortical structures, whose respective grey matter nuclei send and receive neural projections to these regions of the PFC may contribute to this role^23–25^.

Located deep within the brain, the thalamus contributes to multiple sensory, motor and cognitive processes. The ‘first-order’, primary sensory ‘relay’ nuclei, include the ventroposterolateral (VPL) nucleus, which receives sensory afferents from the periphery, and the motor, ventral intermediate nucleus (VIM), which receives afferents from the periphery and cerebellum. The ‘higher-order’ cognitive nuclei are defined by their reciprocal connections with the PFC and a separate group of subcortical nuclei known as the basal ganglia^26^. Higher-order thalamic nuclei include the mediodorsal nucleus (MD), the centromedian nucleus (CM), the centrolateral nucleus (CL), the anterior nucleus of the thalamus (ATN) and the intralaminar parafasicular (Pf) complex^26^. All higher-order nuclei, receive driving input from cortical layer V reflecting a common functional contribution to cognition through enhancement of cortico-thalamo-cortical communication. Nevertheless, cognitive functions may not be exclusively the domain of the ‘higher-order’ nuclei, as lesions of the Ventrolateral anterior (VLa), whose connectivity with the premotor cortex^27^ is more comparable to a first order “relay” nucleus, cause impaired attentional biasing of working memory^28^.

Of the higher-order thalamic nuclei, the functional contribution of the MD nucleus to cognition has been most extensively characterised^29–32^. Its importance in cognition is reflected in dense connections with all frontal cortical regions including the Frontal Pole (FP), the OFC, the vmPFC, the DLPFC, the ACC and the premotor cortex^33^. Accordingly, lesions of the MD nucleus result in behaviour that show striking similarity to those of PFC lesions such as diminished behavioural flexibility, perseveration, and deficits in working memory^34,35^.

The MD nucleus can be divided into sub-regions based upon their connectivity^36,37^. The lateral portion, or parvocellular (MDpc), exhibits strong reciprocal connectivity with the DLPFC (Brodmann areas 9/46) and FP(Brodmann area 10). Consistent with the role these cortical region plays in the formation of abstract rules, patients with lesions of the lateral MD nucleus are impaired in tasks that rely upon rapid learning and re-learning stimulus-action relationships such as the Wisconsin Card Sorting Test (WCST)^38^. In mice, enhancing the activity of the lateral MD nucleus promotes the formation of the cortical neural assembles that form in PFC during stimulus-response learning^39,40^. This has led to the idea that MD nucleus may act to amplify connectivity within the PFC and in doing so enhance the behaviourally relevant rule formation necessary for adaptive-decision making^32,41^. Lesions of the medial magnellocellular (MDmc) subdivision, lead to an in-ability to persist in choosing the most rewarding beneficial decision after it has been identified^42^. The MDmc shares extensive reciprocal connectivity with the OFC^36,37^. The MDmc’s role within this thalamo-cortical network, may be to regulate the threshold of whether to persist, or to change decision strategy, based upon information about a decision’s reward value it receives encoded by the PFC^43,44^.

MR guided Focused Ultrasound (MRgFUS) ablation of the VIM nucleus of the thalamus is routinely performed for patients with medication refractory Essential Tremor^45^. The VIM lies within a central position of the thalamus in close proximity to the higher order thalamic nuclei, including the MD, the CM, but also to the VLa^46^. A characteristic of MRgFUS VIM-thalamotomy is extensive vasogenic oedema, the volume of which increases 4-fold at 24 hours relative to that seen immediately post treatment (Fig.1B). This leads to swelling of both the VIM and its surrounding nuclei ^47^. MRgFUS thalamotomy therefore affords the unique opportunity to prospectively study the lesional effect on the higher-order thalamic nuclei function. By testing patients before and after thalamotomy using the 4­armed “restless bandit” task we aimed to elucidate the thalamic functional role in reinforcement learning, as successful performance in this task relies upon both recruitment of cognitive control and valuation functions of the PFC^8,9^. We hypothesised that if the cognitive thalamic nuclei such as the MD are critical hubs regulating the PFC’s cognitive control or valuation network functions, then oedema extending into this network would produce a behavioural correlate which indexes a loss of cognitive control, value-based learning, or a combination of both.

**Fig. 1.**
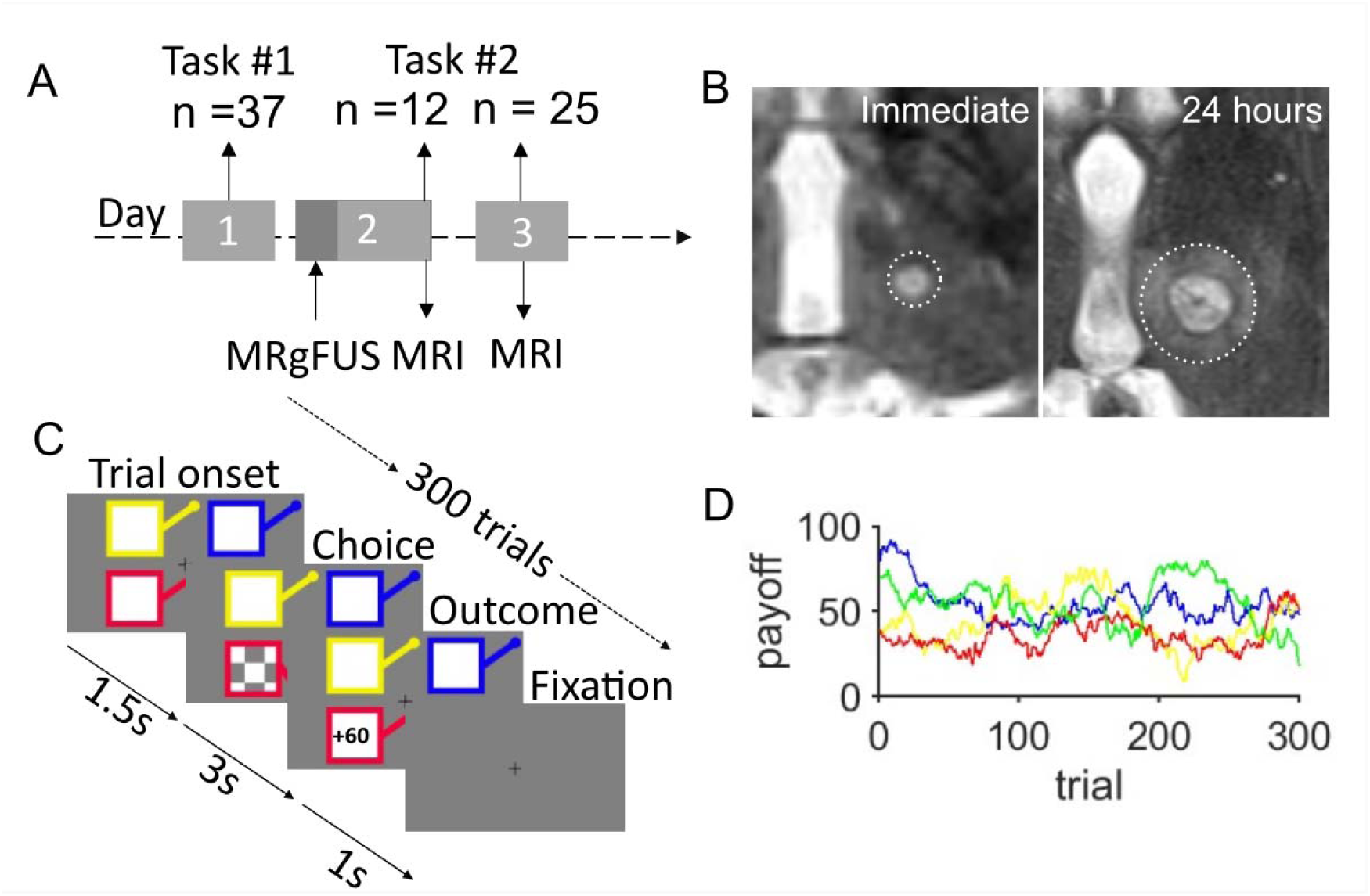
Study design and Restless bandit task. (**A**) Patients performed the task before (n=37) and after focused ultrasound (MRgFUS) thalamotomy. Patients treated in Liverpool (n=12) Post-Thalamotomy performed the task within hours of their procedure. In Dundee (n=25), patients performed the task at ∼24 hours post-op, coinciding with mature post-operative vasogenic oedema (**B**) Representative examples of post-operative thalamotomy in a Liverpool patient scanned within hours of their procedure, and in a Dundee patient where MRI was acquired 24 hours. The different extent of vasogenic oedema is outlined by the white dotted circles. (**C**) Restless bandit task and trial structure. Each trial has a fixed duration of 6 seconds, with a variable intertrial interval (mean 2 seconds). At trial onset, four coloured squares (bandits) were presented. If the participant selected a bandit within 1.5 s, that selection was highlighted by the bandit lever depressing and a chequer board appearing (choice screen). After a 3 s delay the outcome of the choice, as number of “points” won, was displayed for 1 s. In trials where the participant failed to make a choice within 1.5 s, the choice screen was replaced by a large red cross (not shown) signifying a missed trial. (**D**) Example of the underlying payout (reward) structure across the 300 trials of the task for each of the 4 bandits. The payout varied from one trial to the next by a gaussian walk^8^.

Using lesion-behaviour mapping analysis, we show that post-operative lesional oedema extending into the lateral portion of the MD nucleus explains post-operative impairment in choice switching. A reinforcement learning model fitted to this behaviour, suggests the impairment stems from heightened reliance in learned value-estimates which is expressed behaviourally as diminished decision flexibility. These results support a role for the MD nucleus in augmenting cognitive flexibility for adaptive decision making and indicate its potential as a neuromodulation target for disorders of motivation and cognitive control.

## Results

### Behavioural & computational modelling results

Our experimental design tested two mutually exclusive hypotheses: First, if the higher order thalamic nuclei are critical to maintaining cognitive flexibility, then thalamotomy associated oedema should disrupt this function by deafferenting cortical input from and output to the thalamus. Second, we wanted to understand the mechanisms by which the thalamus influences flexible decision making. Does the thalamus regulate the extent to which learnt value estimates are integrated into behaviour? Or alternatively, does the thalamus regulate decision strategies, such as the decision to perseverate, irrespective of the reward obtained?

By testing patients with a task that relies upon learning from, and encoding of, rewarding feedback in the form of a neural value estimate, we could examine the extent to which the thalamus regulates reward sensitivity. This second hypothesis was further enriched by fitting reinforcement learning model variants^9^ which can distinguish between a pure influence on reward sensitivity rather than cognitive inflexibility mediated by perseveration.

With this motivation in mind, we recruited and tested thirty-seven patients with Essential Tremor (ET), in a prospective, case-control design (Fig.1A). Performance was compared to a group of thirty-two age-matched healthy controls (Table 1). The 4-armed restless bandit task was chosen to test of our hypothesis, as the payouts (Fig.1D) for each bandit independently fluctuates (via a Gaussian walk) on a trial-to-trial basis, creating significant choice uncertainty. Choosing the best of the four bandits relies upon a binary choice on each trial, either i) staying with the same bandit, or ii) switching to one of the three alternative bandits. Human behaviour in this task has been robustly reproduced by reinforcement learning models^8,9,48^. These generate trial-by-trial estimates of the value (and uncertainty in value) of each bandit, upon which the PFC has been proposed to guide choices^4^. We therefore reasoned that fitting reinforcement learning models to both Pre- and Post-Thalamotomy behaviour in the task would allow us to examine the influence of the thalamus on value-based decision making.

**Table 1.**
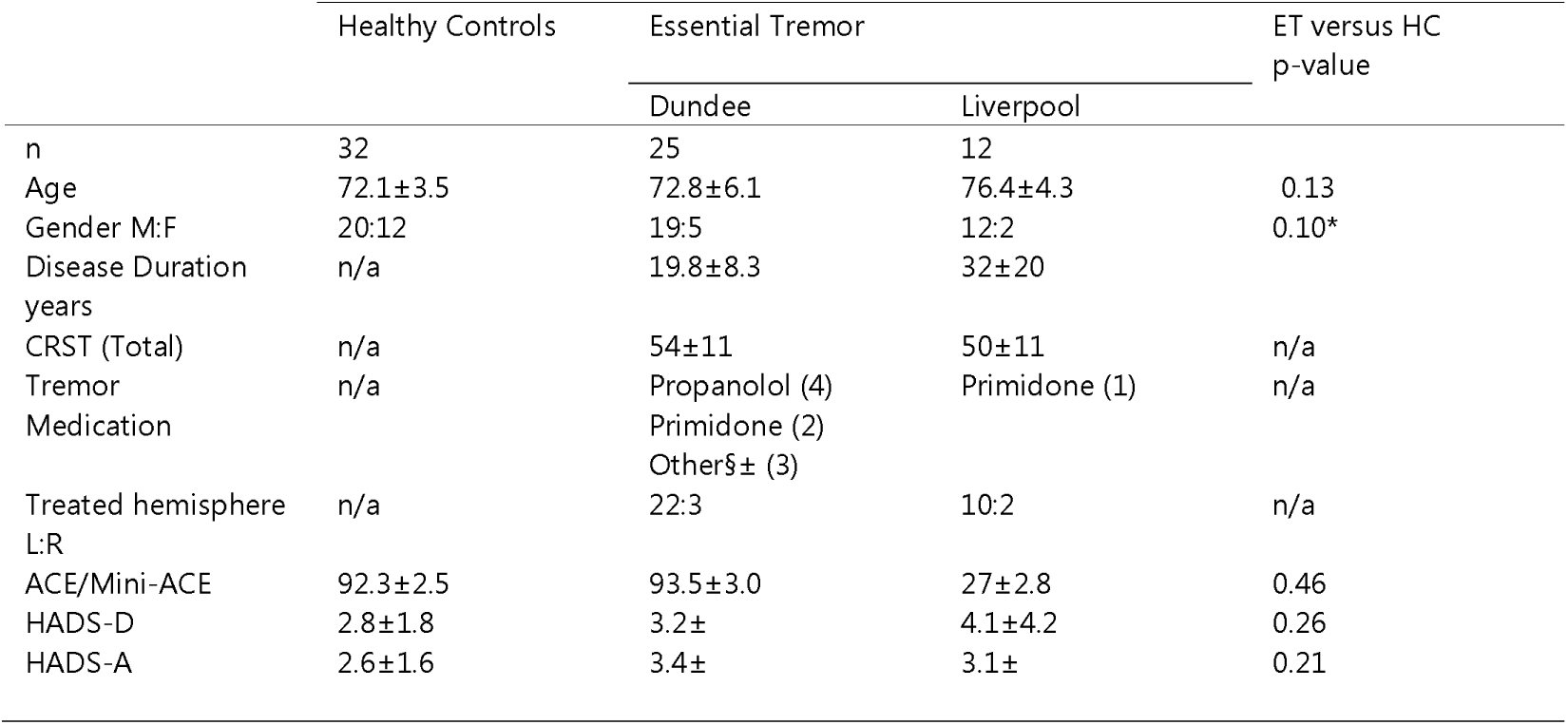
Patient demographics and clinical details. Values expressed as mean ± standard deviation. ET, Essential Tremor, CRST, Clinical Rating Scale for Tremor, HADS, Hospital Anxiety and Depression Rating Scale, MoCA, ACE, Addenbrookes Cognitive Examination. All p-values calculated using Two-Tailed unpaired T-test, Fisher’s exact test. § Other medications included Clonazepam (n = 2) Zonisamide (n = 1).

Each patient performed the 4-armed restless bandit task (Fig.1C) 24 hours before their thalamotomy. Liverpool patients (n = 12) were tested immediately after their treatment (mean 2.5 ± 0.43 hours), while the Dundee patients (n = 25) were tested the day after their thalamotomy (mean 24 ± 0.25 hours). This is due to differences in clinical workflow across sites and allowed elucidation of the relative contributions of non-specific effects of the operation versus the delayed, focal effects of vasogenic oedema on the thalamus (Fig.1B).

Patients identified the best bandit to similar levels as an age-matched healthy control (HC) group (HC = 0.65 ± 0.02), both before and after thalamotomy (Pre-Thal= 0.64 ± 0.03, Post-Thal = 0.62 ± 0.03, effect of group: *F*_1,630_ = 0.09, *p*=0.768, SFig1. A&B). Task performance was unaffected by non-specific effects of the condition or procedure (e.g. diminished attention), as the response times before and after (Pre-Thal = 0.67 ± 0.02, Post-Thal = 0.65 ± 0.02 s) were not prolonged (ANOVA: *F*_1,630_ = 1.02, *p* = 0.35) compared to the control group (HC = 0.66 ± 0.02 s), nor was there a difference between the number of missed trials before or after (Pre-Thal = 3.7 ± 0.5 %, Post-Thal = 3.5 ± 0.6 %), both of which were comparable to the control omission rate (HC = 3.2 ± 0.6 %, ANOVA: *F*_1,630_ = 0.002, *p* = 0.96, SFig1.C&D).

We further examined the effects of thalamotomy on flexibility of decisions by classifying these into either stay or switch choices. An additional fixed effect of procedure site (Liverpool versus Dundee) was included to distinguish between non-specific effects of the procedure over the direct influence of post-procedural oedema within the thalamus. Overall, in the patients, the probability of choosing the same bandit on two consecutive trials, across the task, *P(*Stay), increased after thalamotomy (Pre­Thal = Post-Thal = 0.75 ± 0.02, Pre-Thal = 0.67 ± 0.03, effect of Group ANOVA: *F*_1,437_ = 30.1, *p*<0.001). However, this was driven by the delayed effects of oedema in the Dundee cohort as the Liverpool patients, *P*(Stay) remained unchanged from pre-operative performance (Pre-Thal_Liverpool_= 0.66 ± 0.08, Post-Thal_Liverpool_ = 0.69 ± 0.08). In contrast, patients tested in Dundee 24 hours after their procedure increased their P(Stay) values (Pre-Thal_Dundee_ = 0.67 ± 0.03, Post-Thal_Dundee_ = 0.78 ± 0.04, Group × Site interaction ANOVA: *F*_1,437_ = 7.6, *p*=0.005, Fig.2E). Post-hoc analysis using the linear contrast of the fixed-effects estimates confirmed this was driven by a significant effect of thalamotomy in Dundee compared to their pre-operative performance (*F*_1,437_ = 40.1, *p* <0.001), and compared to the Liverpool patients (*F*_1,437_ = 10.2, *p* = 0.005), where no change in P(Stay) was observed (*F*_1,437_ = 1.11, *p* = 0.5). Increased stay choices, Post-Thalamotomy, could not be explained by simple lateralisation bias of a unilateral lesion, as the proportion of choices made to the two bandits on the left and right of the presentation screen (0.44 ± 0.04) was no different in the Dundee patients than before their thalamotomy (0.42±0.05) (effect of Group ANOVA: *F*_1,437_ = 2.3, *p* = 0.18; Group × Site interaction ANOVA: *F*_1,437_ = 2.8,*p* =0.11). Non-specific cognitive effects of the thalamotomy could not explain this change in decision flexibility as there was no difference in the overall task performance, indexed by the proportion of choosing the best bandit (Group × Site interaction ANOVA: *F*_1,437_ = 0.93, *p* = 0.33, Fig.2A&B), proportion of missed trials (Group × Site interaction ANOVA: *F*_1,437_ = 0.03, *p* = 0.85, Fig.2F). Arguing against diminished attention leading to choice perseveration, the patients in the Dundee cohort were faster (Pre-Thal_Dundee_ = 0.69 ± 0.03 s, Post-Thal_Dundee_ = 0.65 ± 0.03 s) than the Liverpool patients (Pre-Thal_Liverpool_ = 0.64 ± 0.04 s, Post-Thal_Liverpool_ = 0.65 ± 0.05 s) to make their decisions (Group × Site interaction ANOVA: *F*_1,437_ = 10.01 *p*=0.001, Fig.2C), a between-group effect that was mediated by the difference in the Dundee patients pre and Post-Thalamotomy(*F*_1,437_ = 9.2, *p* = 0.01). A difference in the reward payout received by the different patients could not explain the difference in behaviour as there was no significant difference in the payoff schedule allocated to each patient pre-and Post-Thalamotomy (/2 37 = 0.64, *P* = 0.72).

**Fig. 2.**
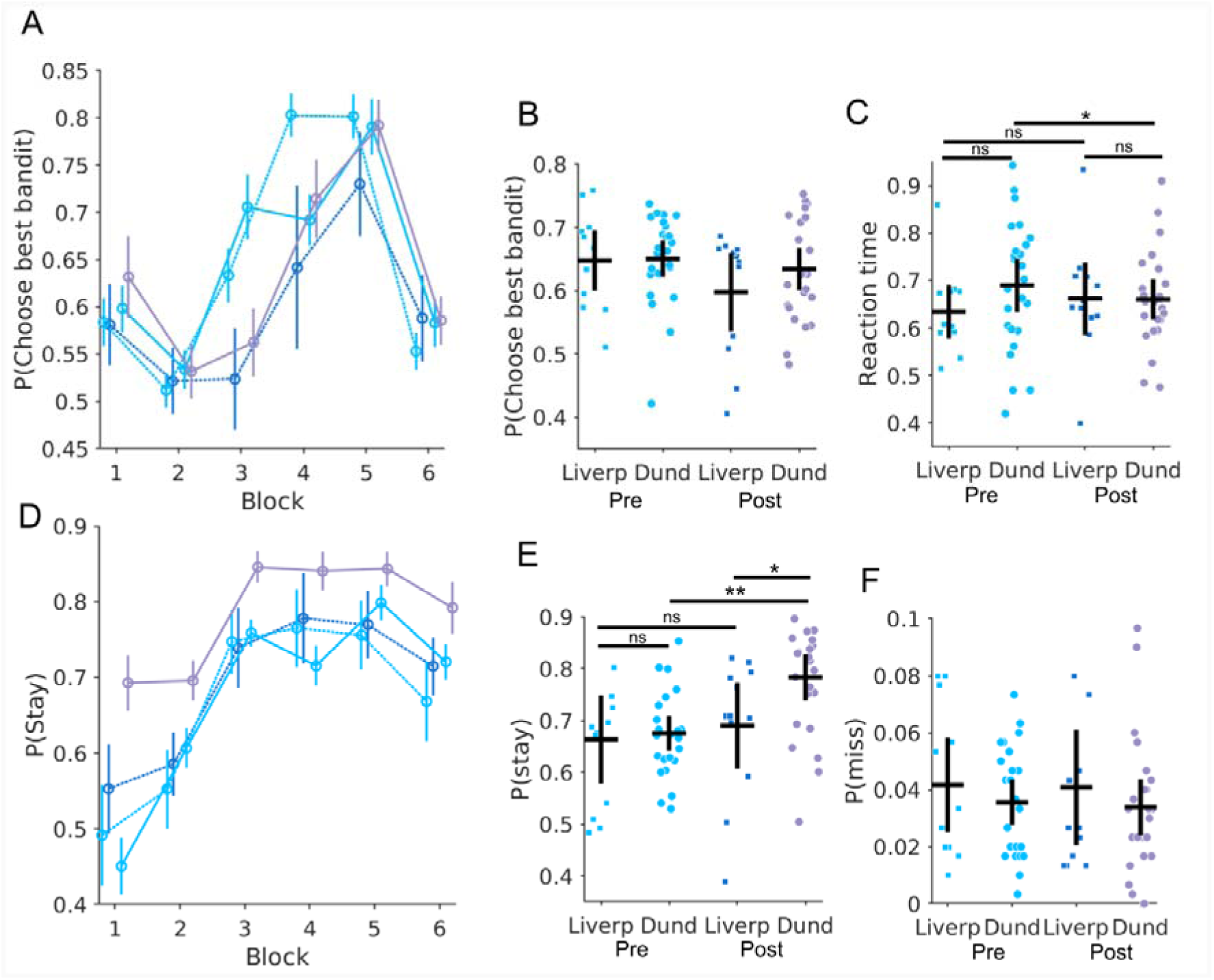
Delayed effects of Thalamotomy on flexibility of decision making in the restless bandit task. The average probability of choosing the bandit with the highest payout, P(Choose best bandit) is plotted pre (light blue) and Post-Thalamotomy (dark blue). There was no effect of the thalamotomy oedema on the ability of the Dundee patients to identify the best bandit. Solid lines represent performance in patients tested 24 hours after thalamotomy (Dundee cohort n = 25), dotted lines represent patients tested immediately following (∼1-2 hours) thalamotomy (Liverpool cohort n = 12). (**A**) The increase in average values of best bandit choice plotted across all six blocks of 50 trials. Vertical lines S.E.M. In (**B**) each circle represents average best bandit choice probability across the task for an individual subject from the Dundee cohort (squares represent the Liverpool cohort). The horizontal and vertical bar represents the group mean and 95% confidence limits. (**D**) Probability of stay choices P(Stay) across six bins of 50 trial, plotted for pre & Post-Thalamotomy for both groups. (**E**) Individual average P(Stay) values across the whole task, *p=0.005, **p<0.001. Average reaction time between groups across the task *p=0.01 (**C**) and proportion of missed trials (**F**) for each group. There was no cross-site effect on the proportion of missed trials. All p-values reported are Bonferroni corrected. ns = not significance difference.

Staying with the same option, rather than switching, could be explained by two mechanisms. One may be a pure loss of cognitive flexibility, whereby adaptive, stimulus-response mapping is replaced with a sub-optimal strategy of choice perseveration. Alternatively, choice repetition could be driven by exploitative decisions which aim to maximise the reward obtained by choosing the options with the highest estimated value, at the expense of explorative, evidence gathering behaviour. Computational models can be used to distinguish between mechanisms by varying how the models learnt value estimate is incorporated into the model’s choice rule. For example, perseveration can be modelled by a perseveration bonus which multiplies a *single* actions value estimate. In contrast, a reward sensitivity parameter, which multiplies the influences of *all* actions learnt value estimate, can delineate the extent to which choice repetition underpins value exploitation. We fitted a series of previously validated model variants ^13,14^ to the patient’s choices which include, as free parameters, reward sensitivity, an exploration bonus and a perseveration bonus. By modelling the value and uncertainty in value of each of the four options throughout the task, we classified stay/switch choices further, with a trinary categorisation based upon the model’s value estimate.

An exploitative choice refers to the selection of the option perceived to have the highest expected value. A directed exploratory choice is made towards one of the three less valued options, particularly the one with the highest level of uncertainty (which in general, has been least recently selected)^15^. Directed exploration scales linearly with the modelled exploration bonus. Random exploratory choices involve the selection of a less valued option regardless of current knowledge or uncertainty about their payout^49^. In contrast to directed exploration, which actively seeks information to update the estimate of a choices value, randomly explored options reflect indifference to the value-estimate and can be linked to decision noise^50^. The effect of thalamotomy on these choice types was analysed by estimating each subject’s proportion of each choice through the task. The probability of making an exploitative, random or directed exploratory choice were designated as P(Exploit), P(RE) and P(DE) respectively.

Leave one out (LOO) cross-validation estimates^51^ were computed for each model variant and to pre­post-operative patient and healthy control groups. Consistent with prior studies^9,48^, the Bayesian Learner model with terms for directed exploration and perseveration bonuses (Bayes-SMEP) was the model which most accurately represented all groups, including Pre-Thalamotomy (loo-log likelihood_Pre_ = −0.687) and control groups (loo-log likelihood_HC_= −0.740), Fig. 3A & Supplementary Table 1. Following thalamotomy, though the Bayes-SMEP continued to be the winning model, there was a marked improvement the model fit in the Dundee group (loo-log likelihood_Dundee_ = −0.458), which was not observed in the Liverpool patients (loo-log likelihood_Liverpool_= −0.663). This improvement in model fit in the Dundee cohort was seen across all eight model variants (Fig. 3A & Supplementary Table 1) and was independent of the model comparison metric employed (Supplementary Table 1). Consistent with the winning model capturing decision making in the task robustly, model parameters could still be recovered (Fig.3 B-D) from synthetic choice data generated from simulated choices. Simulated choice data from the winning model also overlapped with the experimental choices, replicating both the proportion of experimentally observed *P*(Choose best bandit), and *P*(Stay) choices, (Fig. 3 E-H) across each of the 6 analysis blocks of the task, and across the task as a whole.

**Fig 3.**
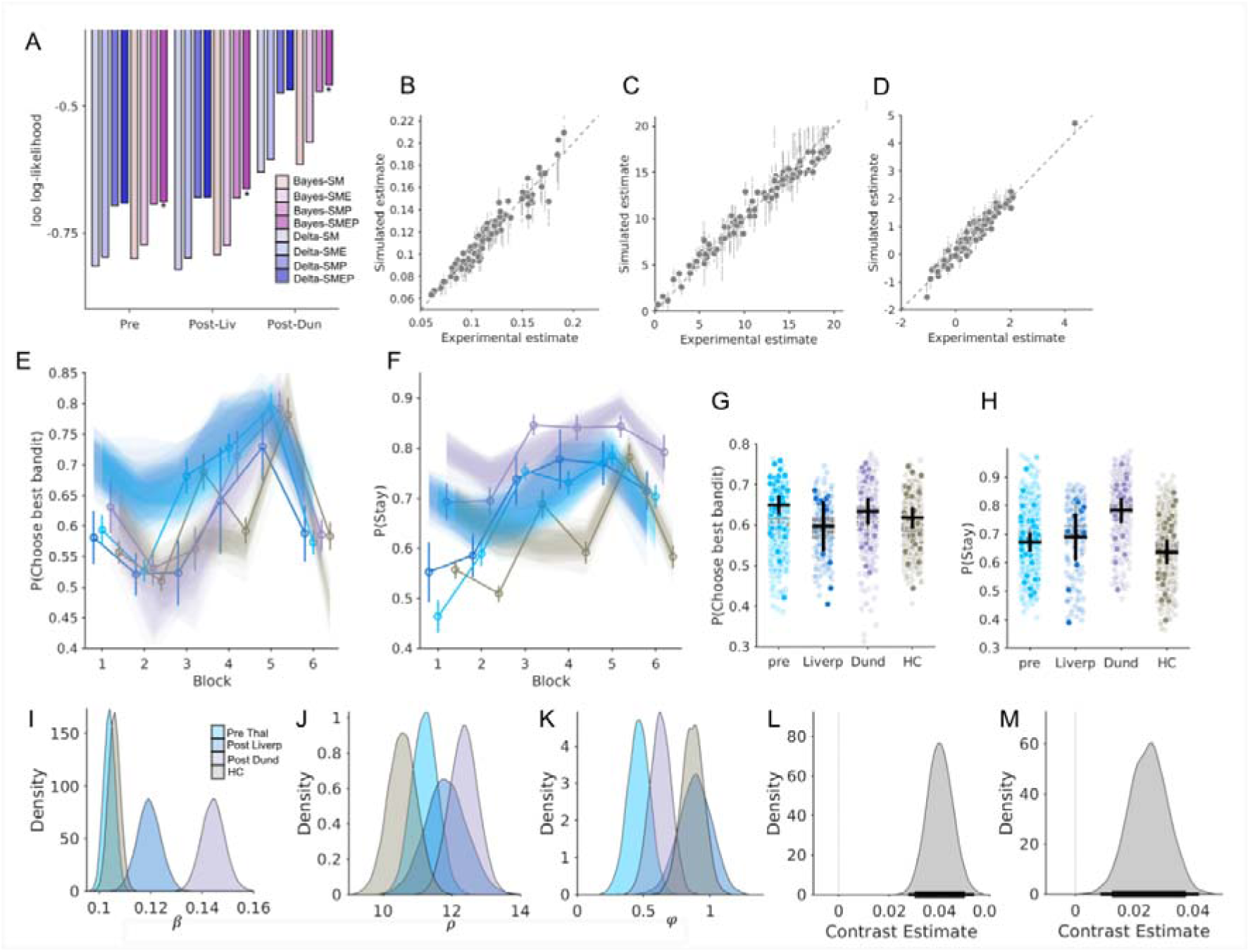
**Reinforcement learning model fitting to thalamotomy choice reveals mechanism of cognitive inflexibility.** (**A**) Model fit (Bayes/ Delta learning rules with 4 choice rule variants) fitted to Pre-Thalamotomy, and post-thalamotomy for both the Liverpool (Liv) and Dundee (Dun) groups. Each group’s winning model (Bayes-SMEP) delineated by *. (**B-D**) Each grey dot represents average parameter estimate recovered from fitting 15 experiments simulated using each individual’s estimated parameter fit. The vertical lines represent the 95% higher density interval (HDI). Successful parameter recovery for the reward sensitivity parameter, (B), perseveration bonus,,(**C**) and exploration bonus,, (**D**) was indexed by in strong linear correlations between the experimentally estimated parameters and those recovered from simulated choices (; R^2^ = 0.93, ; R^2^= 0.97, ; R^2^ =0.93, all p<0.001). In (**E & F**) the experimental best bandit choices P(Choose best bandit) and stay choices, P(Stay) are plotted, across six blocks of 50 trials. The same analysis is applied to the simulated model choices for each group. Shaded areas present 95% confidence limits across simulations. (G) Simulated individual P(Choose best bandit) average are in light coloured dots, overlayed with experimental equivalent with dark coloured. Black vertical and horizontal lines are experimental mean and 95% confidence limits, grey dotted lines simulated. (H) Illustration of simulated versus experimental P(Stay) values through the task demonstrating the simulated data could replicate both the increase in these choices Post-Thalamotomy in the Dundee and absence of this effect in the Liverpool patients. (I) Posterior distributions for, (I) and, (J) and (K) illustrated for each patient group and healthy controls (HC). (L) Posterior differences in the mean for, Contrast: Post-Thal Dundee minus Pre-Thal and in (M) Contrast: Post-Thal Dundee minus Post-Thal Liverpool. Thin and thick horizonal black lines represent 95 and 85% HDI.

Post-Thalamotomy, both groups of patients had significantly higher values of the reward sensitivity parameter, *β* (posterior difference in mean (contrast: Pre-Thal minus Post-Thal, Dundee M_diff_ = 0.04 (HDI = [0.03,0.05]), Liverpool M_df_ = 0.01 (HDI = [0.002,0.03])). The increase in *β* was significantly greater in the Dundee cohort (contrast: Post-Dundee minus Post-Liverpool, M_diff_ = 0.02 (HDI = [0.01,0.04]), Fig.3 I,L-M). No difference in the perseveration bonus, p, was observed in either group (Dundee M_diff_ = 1.13 (HDI = [-0.26, 2.26]), Liverpool M_diff_ = 0.62 (HDI = [-1.24, 2.46]), Supplementary Table 2). The exploration bonus was unaffected by the thalamotomy (Supplementary Table 2) but was found to be significantly lower in pre-operative ET patients than the healthy controls (contrast: Pre-all minus healthy control, φ,M_dff_= −0.40 (HDI = [−0.72, −0.10]).

Using the value and uncertainty estimates from the winning model (Bayes-SMEP) to divide each choice into the trinary classification described above, the Dundee patients made more exploitative, *P*(Exploit) choices through the task after thalamotomy than the Liverpool group (Post-Dun = 0.79 ± 0.03, Post-Liv = 0.72 ± 0.05) who made a similar proportion Pre-Thalamotomy (Pre = 0.73 ± 0.02, Fig.4 A,D, Group × Site interaction ANOVA: *F*_1,437_ = 3.83, *p* = 0.05). Using the linear contrast of the fixed-effects estimates, this interaction was driven by a significant increase in *P*(Exploit) in the Dundee patients after thalamotomy compared to both their own pre-operative choices, and the Liverpool patients post-operative choices (Dundee: *F*_1,437_ = 9.6, *p* <0.01, Liverpool: *F*_1,437_ = 6.6, *p* = 0.04). Thalamotomy in the Dundee cohort correspondingly reduced the *P*(RE) post-operatively compared to both the Liverpool patients and pre-operatively (Dundee = 0.13 ± 0.02, Liverpool = 0.17 ± 0.03, Pre = 0.17 ± 0.01), Fig. 4, B & E, Group × Site interaction ANOVA: *F*_1,437_ = 4.82, p = 0.028). Again, this interaction was mediated by the reduction in P(RE) in the Dundee patients compared to their pre-operative performance (*F1,437* = 13.8, *p*<0.001). Consistent with reduced *φ* value above, as exploration bonus linearly scales with the proportion of directed exploration P(DE), Pre-Thalamotomy P(DE) was less than in healthy controls (HC = 0.11±0.01, Pre-Thal = 0.09±0.01, effect of group ANOVA *F*_1,437_ = 5.6, p = 0.017). However, pre-operatively neither P(Exploit) or P(RE) were different to the HC group (P(Exploit) effect of group *F*_1,437_ = 1.2, *p* = 0.26, P(RE) effect of group *F*_1,437_ = 0.25 *p* = 0.61). The effect of thalamotomy in the Dundee patients further suppressed P(DE) through the task, a reduction that was not observed in the Liverpool patients (Dundee = 0.06±0.01, Liverpool = 0.11±0.02, Group × Site interaction ANOVA: *F*_1,437_ = 7.21, p < 0.01, Fig. 4 C&F). Analysing the between-group differences confirmed that this was due to the reduction in the Dundee patients compared to their pre-operative performance (*F*_1,437_ = 16.7, p<0.001) and the difference between the Liverpool and Dundee patients Post-Thalamotomy (*F*_1,437_ = 17.58, p<0.001).

**Fig 4.**
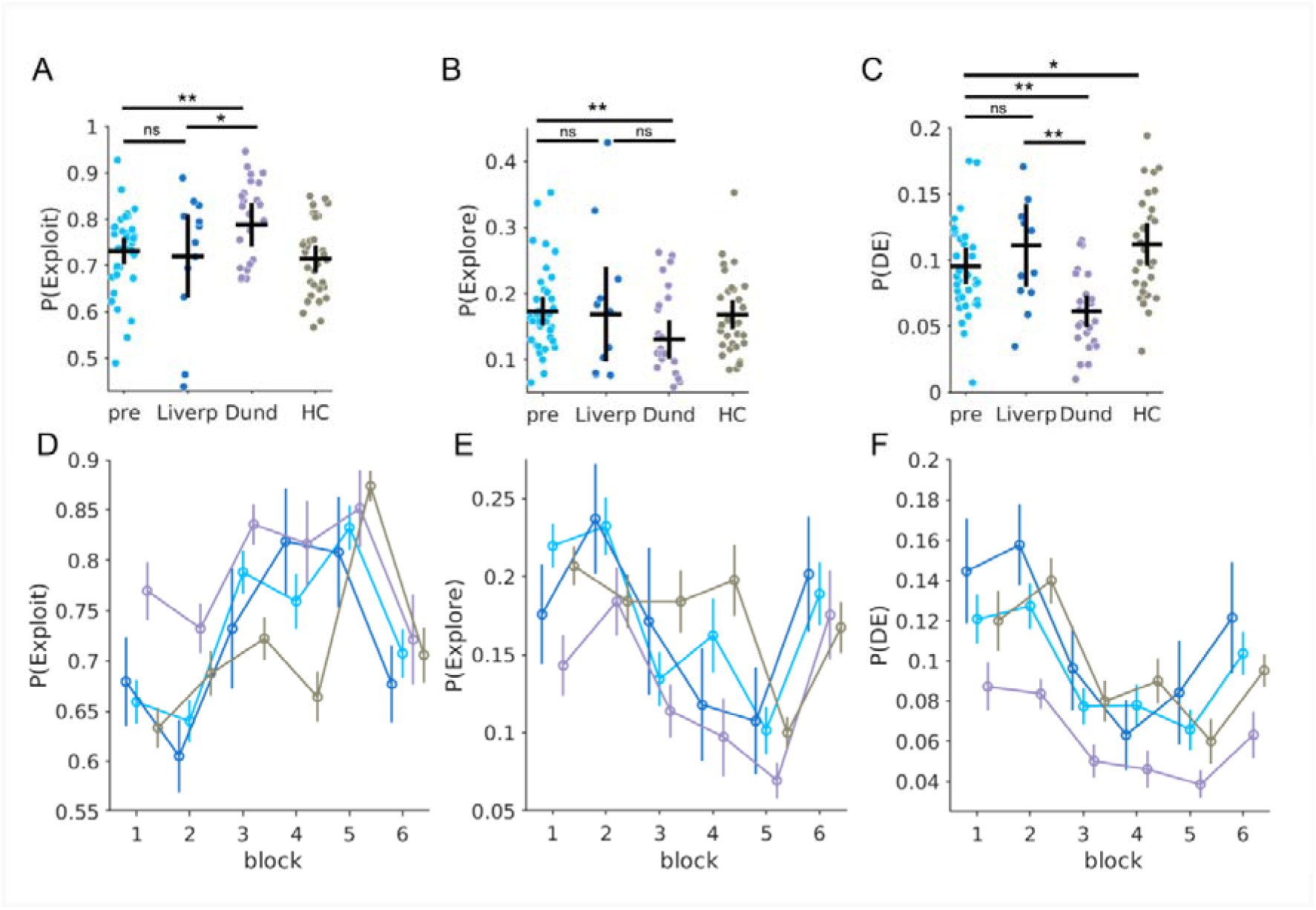
**Effect of thalamotomy on value-based decision types. (A)** Proportion of choices to the highest value option P(Exploit) across the task in the four groups.\**p*< 0.05,** *p*<0.01 (**B**) Proportion of random exploratory choices, P(Explore), ** p<0.001, and (**C**) directed exploratory choices p(DE) \*\**p*<0.001, \**p* <0.05. Each dot represents an individual patient with Pre-Thalamotomy in light blue dots, Post-Thalamotomy (Liverpool) in dark blue dots, Post-Thalamotomy (Dundee) in purple dots, and healthy controls (HC) are in grey. Black horizontal and vertical lines represent group mean and 95% confidence limits. Other than those significant differences delineated by an asterix no other differences between groups were significant. The time course of each decision type across six blocks of 50 trials for all groups is plotted in D to F. *P*-values are Bonferroni corrected. ns = not significant.

These results confirmed that the increase in stay choices Post-Thalamotomy was best explained by several factors. Firstly, reflected by the increase in /?, choice repetition was driven by a greater extent to which decisions were biased by the neural estimate of all four bandits value. This was expressed into the task behaviour by a switch towards more exploitative choices to the higher valued option, and reduced random section of lower valued alternatives. Secondly, the reluctance to switch choices can also be explained by an increase in reliance upon value estimates, as directed exploration - a decision strategy which is aimed at updating the value estimate of bandits with high uncertainty, was significantly suppressed by thalamotomy.^49^

### Neuroimaging correlates of thalamotomy induced cognitive inflexibility

Next, in the Dundee cohort, we analysed the relationship between the extent of post-operative vasogenic oedema into individual thalamic nuclei and the main behavioural effect of thalamotomy, increased stay choices ΔP(Stay) (P(Stay)_Post_ – P(Stay)_Pre_). The extent of post-op oedema within the thalamus as a whole did not predict the increase in stay choices (mean oedematous voxels = 2102 ± 241, rho(23) = −0.31, Bonferroni adjusted *p* = 0.1). Furthermore, estimates derived from individualised segmentations of the thalamic nuclei which were encapsulated by the oedema (VLa, MD, VIM, CM, Fig. 5A) did not correlate with ΔP(Stay) (Supplementary Table 3 & Supplementary Fig. 2). We therefore reasoned that this effect could be mediated by the pattern of oedema extension at a finer spatial resolution. To test this hypothesis, we fitted a voxel-wise GLM^52,53^ to identify a thalamic “sweetspot” which predicted ΔP(Stay) before and after thalamotomy. Applying this approach to voxels where thalamotomy oedema consistently involved the MD nucleus (Fig 5B) revealed a cluster of voxels where the presence of oedema positively correlated with the increase in ΔP(Stay) (Peak voxel GLM beta weight = 2.75, MNI (*x, y, z)*: (−8,−19,6); voxel permutation test *p* = 0.01, cluster permutation test *p* = 0.01, Fig.5 C&D, Supplementary Table 4). No other clusters where thalamotomy oedema correlated positively with increased ΔP(Stay) survived permutation testing when this analysis was applied to the VLa, VIM, CM (Supplementary Fig. 3 & Supplementary Table 4). We also found evidence that the distribution of oedema within the thalamus was mediating Post-Thalamotomy choice inflexibility as clusters which negatively correlated with the ΔP(Stay) were identified within the MD (beta weight = - 2.6, MNI (*x, y, z)*: (−7,−21,−1), voxel permutation test *p* = 0.002, cluster permutation test *p* = 0.004, & beta weight = - 2.13, MNI (*x, y, z)*: (−8,−21,5); voxel permutation test *p* = 0.01, cluster permutation test *p* = 0.005, Fig.5 C&D) but also within the neighbouring CM (beta weight = - 2.59, MNI (*x, y, z*): (−10,−24, 1), voxel permutation test *p* = 0.01, cluster permutation test p = 0.01) and VLa (beta weight = −2.33, MNI (x,y,z) : (−13,−11, 1), voxel permutation test *p* = 0.03, cluster permutation test *p* = 0.02, Supplementary Fig. 3 & Supplementary Table 4).

**Fig 5.**
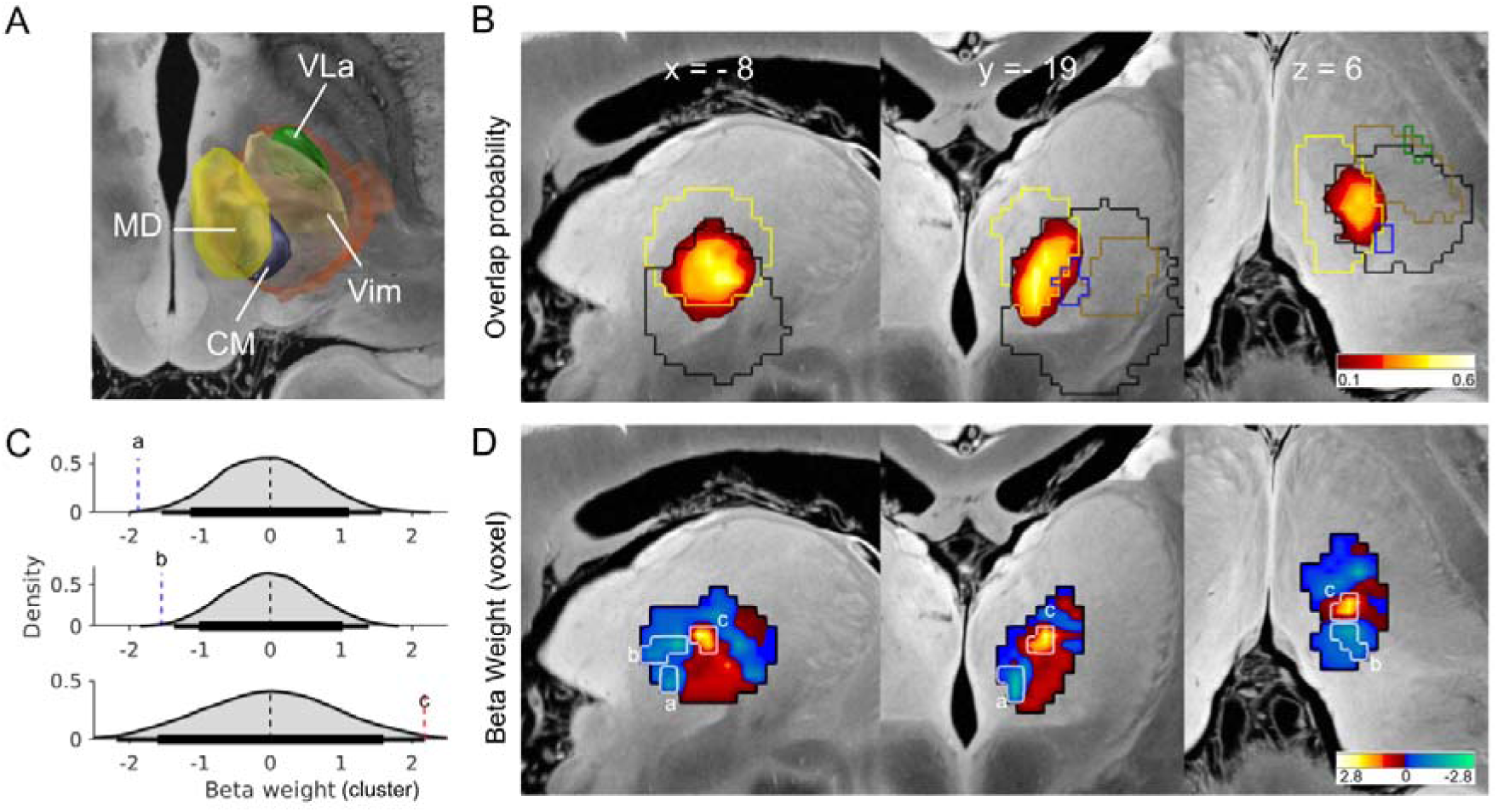
**Spatial distribution of thalamotomy oedema correlates with changes in behaviour.** (**A**) 3D projections of thalamotomy oedema overlap (orange) across all patients in relation to the Ventral intermediate (VIM, brown) and surrounding centromedian (CM, blue), Ventro-lateral anterior (VLa, green) and Mediodorsal (MD,yellow) nuclei. Background from Edlow et al., (2019)^54^. **(B**) Sagittal, coronal and axial slices with the yellow-red heatmap illustrate the probability of lesion oedema extending into the MD nucleus across 24 patients at 24 hours following thalamotomy in the Dundee patients. The outer border of the MD (yellow), CM, (blue) VIM (brown) and VLa (green) are superimposed alongside the outer oedema margins used in A (**D**) The beta weights from a voxel wise GLM absence of oedema was calculated for each voxel within the MD using a GLM correlating with the individual patients **Δ**P(Stay) over the task, (P(Stay) post-minus Pre-Thalamotomy). Warm coloured Beta values indicate voxels where the presence of oedema is associated with increased P(Stay) cold coloured denote the negative Beta weights which reflect negative correlations. The black outline around the beta weight heatmap represents the outer border of the voxels analysed were at least 75% of patient’s oedema overlapped with the outer border of their MD nucleus following individualised segmentation of the thalamus. Voxel and cluster-based permutation testing confirmed three clusters with both positive and negative predictive value illustrated by the white outer border (a,b,c). Each cluster survived correction for multiple comparisons using two tailed permutation threshold of p<0.05. Cluster level permutation distributions for each cluster our illustrated in (C) horizontal line represent 95% (thick) and 99% (thin) confidence limits.

We calculated the voxel most likely to be associated with the behavioural effect. Weighting each individual patient MD-lesion mask by the change in ΔP(Stay) (P(Stay)_Post_ – P(Stay)_Pre_) confirmed an infero-lateral portion of the MD nucleus was most likely to increase ΔP(Stay) (voxel-Wise Signed Rank Z = 3.8, *p*<0.05, *FDR q* = 0.05, MNI co-ordinate x = −7.9, y = −17, z = 3).

Using the overlap of thalamotomy oedema into each segmented nuclei (MD, VIM, CM and VLa) as seed regions to a group connectome derived from resting state functional connectivity of healthy volunteers^55^, we derived whole-brain functional connectivity maps for each patient (Fig. 6). The correlation between the functional connectivity and the thalamotomy induced ΔP(Stay) was calculated on a voxel-by-voxel basis. The only seed region whose connectivity pattern could predict this effect was the MD nucleus. The R-Map in Fig.6 A depicts thresholded correlations between functional connectivity from thalamotomy oedema volumes to all brain voxels and thalamotomy induced increase in stay choices. Spatial correlation of individual patients’ thalamic seed connectivity profiles with this R-Map significantly explained variants in the between subject increase in P(Stay), (*R* = 0.61, *p* <0.001, permutation test: *R* = 0.65, *p* = 0.004). Supplementary Table 5 includes the details of the peak R-map values and voxel co-ordinates of the cortical regions whose connectivity with the MD nucleus predicted increased stay choices. The same analysis applied to the VIM, CM or VLa did not reveal statistically significant R-maps which exceeded the permutation testing threshold (VIM: *R* = 0.32, *p* = 0.11, permutation test *R* = 0.33, *p* = 0.14; CM: *R* = 0.40, *p* = 0.06, permutation test *R* = 0.38, *p* = 0.09; VLa: *R* = 0.42, *p* = 0.04, permutation test *R* = 0.43, *p* = 0.1, Supplementary Fig. 4).

**Fig 6.**
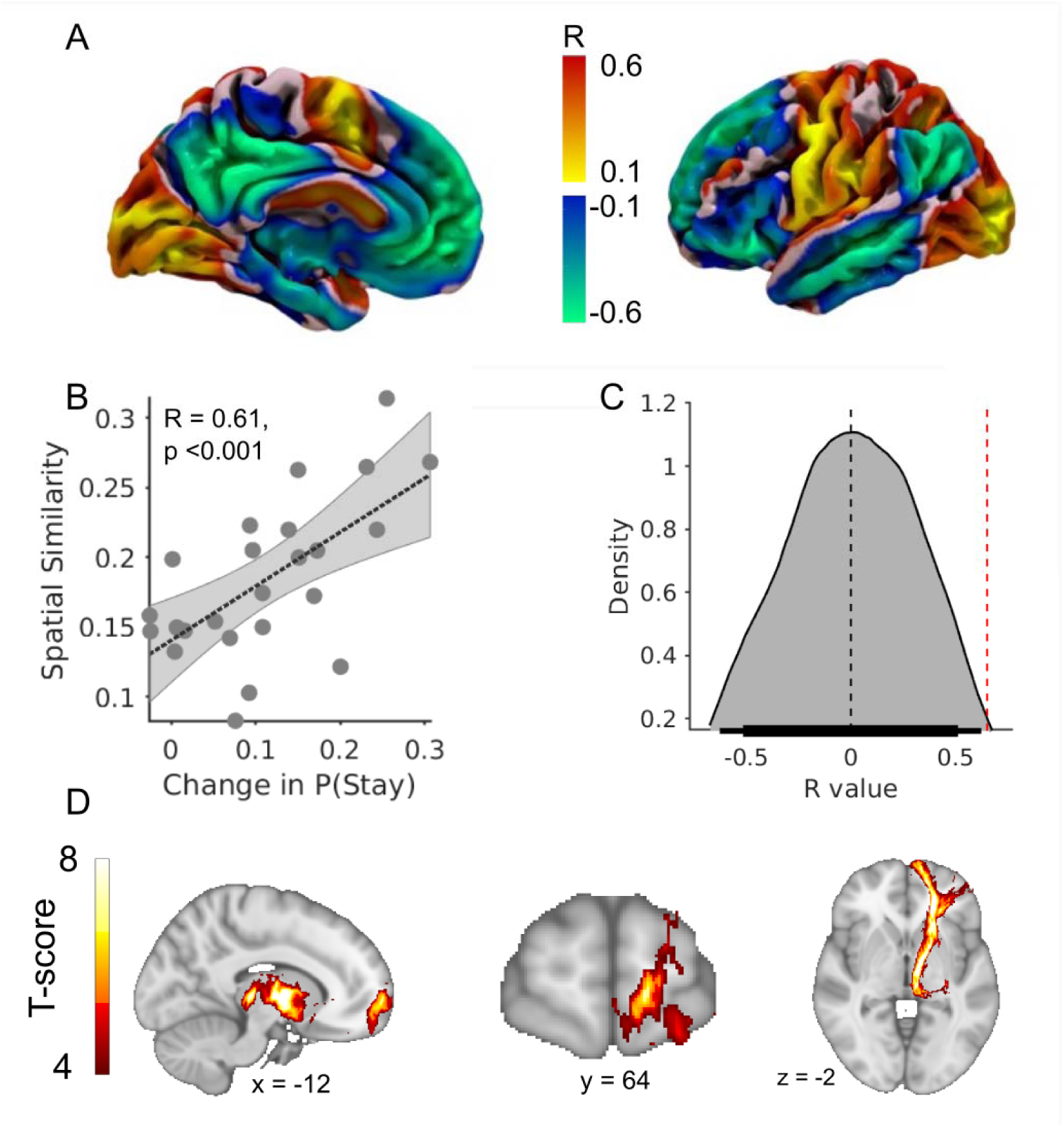
**Mediodorsal thalamus-prefrontal cortical connectivity predicts thalamotomy effect on choice persistence.** Medial (**A**) and lateral cortical surface projections of the whole-brain voxel-wise R-Map demonstrates the optimal functional connectivity profile between thalamotomy oedema extension into the MD nucleus and increase in P(Stay). Warm colours show cortical voxels where functional connectivity to the voxels of the MD nucleus associated with increased P(Stay). Cool colours indicate voxels where stronger functional connectivity was associated with less marked effect of thalamotomy on the same behaviour. **(B**) The more the individual functional connectivity profile matched the ‘optimal’ R-Map, the greater was the increase in P(Stay) induced by thalamotomy related oedema into the MD nucleus (R = 0.61, *p*<0.001). In (**C**) we plot the R value distribution derived from 1000 re-permuted correlations between the individual **Δ**P(Stay) caused by the thalamotomy and the optimal R-map. The probability of seeing the same correlation by chance was *p*<0.01 (represented by the red dashed vertical line). Horizontal thin and thick lines represent 95 and 85% HDI. (**D**) Group level T-scores of structural connectivity between thalamotomy overlap with MD nucleus and the prefrontal cortex peak connectivity in Brodmann area 10, Frontal pole MNI T = T = 7.14 [−16, 57, −10], pFWE <0.001. Colour bar thresholded for all voxels with significant connectivity FWE corrected at p<0.05.

To further validate the conclusion that the increased stay choices were mediated by lesion effects on the MD nucleus to heighten reward sensitivity, we repeated this analysis using the difference in exploitative choices after thalamotomy as the behavioural regressor. As for the analysis of stay choices, the total of oedema volume of the thalamotomy or any other thalamic nucleus including the MD, did not correlate with the increased in proportion of exploitative choices, (rho(23) = −0.11, *p* = 0.6, Supplementary Fig. 2 A-H & Supplementary Table 2). However, increased exploitation caused by the thalamotomy could predicted by the functional connectivity profile of the overlap between the thalamotomy oedema and the MD nucleus with the rest of the brain (*R* = 0.61, *p* < 0.001, permutation test *R* = 0.62, *p* = 0.005, Supplementary Fig. 5 & Supplementary Table 6).

Finally, we performed structural connectivity analysis on individual patient DTI images acquired pre- operatively, to confirm which prefrontal cortical regions shared white matter tract with the region of the mediodorsal nucleus swollen by the thalamotomy. Probabilistic fibre tracking confirmed significant clusters of connectivity between voxels where oedema extended into the MD nucleus and prefrontal cortex which was maximal with the frontal pole (One sample T-test *T* = 7.74, *p*<0.001 TFCE-FWE-corrected; voxels=37884; MNI (*x, y, z)*: (−16,57,−10), Fig.6 D) but in addition exhibited significant clusters of connectivity with the DLPFC (*p*<0.05 TFCE-FWE-corrected, Supplementary Table 7). In a further exploratory analysis, we correlated the change in task behaviour with the thalamo-cortical structural connectivity profile. Significant clusters of connectivity which positively correlated with both the increase in ΔP(Stay) choices and Δ*P*(Exploit) choices following thalamotomy were identified DLPFC (BA 9/46), OFC (BA11) and Frontal Pole (all p<0.05 TFCE-uncorrected), Supplementary Tables 8 & 9.

## Discussion

In this study we examined the immediate behavioural consequences of MRgFUS thalamotomy on the 4-armed restless bandit task. A significant body of research has implicated the prefrontal cortex as the core processing hub where the neural estimate of a decision’s value is integrated into the eventual decision, guiding selection towards the optimal goal^12,41^. Via constant updates from experiential learning, decisions are adjusted to rapidly changing environmental conditions^8^. Prior to this study, whether the thalamus might influence the way decision value estimates are integrated into human behaviour has been proposed but not firmly established^57^. Here we provide causal evidence that an immediate consequence of transiently deafferenting the cortico-thalamic circuit is a marked reduction in decision flexibility. This observation would support a role for the physiologically intact thalamo­cortical circuit in maintenance of choice flexibility, a result supported by clinical observations that medial thalamic lesions typically mimic the effects of those seen in lesions of the prefrontal cortex (such as frontal perseverative syndrome)^58^. A simple explanation such as cognitive inattention or laterality bias could not explain this effect, as our patients were no slower, less accurate, nor more inclined to choose bandits displayed to one hemifield over another, than before thalamotomy. Furthermore, this effect on the task was seen exclusively in the group who performed the task once thalamic oedema had evolved. Despite both patient groups receiving the same procedure, this effect was observed exclusively in those patients tested at a time when thalamic oedema was sufficiently mature to envelop and disrupt the function of the higher order thalamic nuclei. This dissociation between the groups, provides strong evidence that the change in task performance is unlikely to be the consequences of by non-specific effects of the operation. A more likely explanation is that this can be attributed directly to the delayed development of vasogenic oedema produced by the effects of focused ultrasound ablative neurosurgery and the disconnection that this induces between the thalamus and cortex.

By fitting a computational model to the patient’s behaviour, our results further elucidate the specific influence of the thalamus on decision flexibility. We included model variants whose free parameters, reward sensitivity *(β),* exploration, *(φ)* and perseveration, *(>),* bonuses, allowed us to identify how the thalamotomy influenced different aspects of the decision value estimate. Cognitive inflexibility, in the form of response perseveration, as classically described in prefrontal lesions, is unlikely as p, which augments a *single* action’s value estimate to promote choice repetition, was unaffected by thalamotomy. Instead, simulated choices using the winning model (Bayes-SMEP, Fig.3 F&H), could reproduce the effect of thalamotomy only with a significant increase in the reward sensitivity parameter *β.* Because this multiplies the value estimate for *all* actions, decreased decision flexibility is best explained by an overall increase in the patient’s choices being influenced by the learnt value estimates across all options rather than one option in isolation. By this mechanism, our modelling suggests that following thalamotomy, our patients make more stay choices because they have greater reliance upon the neural value estimates to maximise reward seeking. Accordingly, we found a shift towards more exploitative choices and fewer exploratory ones, consistent with heighted value-based decision making mediating an effect on the explore-exploit trade-off^59^. Interestingly, this interpretation is consistent with a proposed role for thalamo-prefrontal coupling in promoting changes in decision strategy under conditions of uncertainty^40,57,60–63^. By decoupling this negative feedback loop, thalamotomy effectively raised the threshold for questioning the need to change decision, leading to a greater likelihood of repeating the same choice. An alternative interpretation is that increased stay choices reflect heighted resistance to environmental volatility. In the intact thalamo-cortical circuit, relatively small differences in payout from one trial to another, or in the relative payout between bandits, lead to a switch in choice. In contrast, post-thalamotomy, these differences need to be greater to promote a switch in bandit. From a modelling perspective, resistance to payout volatility would be expected in a model with a greater *β* parameter, as this drives a deterministic choice policy whereby the softmax output probability maximises choices with large differences in their estimated value.

We also found a significant reduction in the proportion of directed exploratory (DE) choices. Choices which explore lower valued bandits are classified as directed, rather than randomly explored, when these are made to the option which has the greatest choice uncertainty. By actively choosing the bandit with least recent sampling history, the information gained from directed exploration is used to update and reduce uncertainty in the agent’s model, in this case the value estimates, of the environment’s structure. The effect of the thalamotomy to substantially reduce this type of choice, can be considered analogous to diminishing curiosity^64^ or reflective, prospective decision-making^65^ about the reward environment. In combination with an overinflated estimate of the bandit’s value (from the increase in *β),* the behavioural indifference to change leads to maintenance of a decision *status-quo*.

The reduction in DE was paralleled by a striking improvement inthe model fits across all variants (including the winning Bayes-SMEP). Even the simplest model (Alpha-SM) was a better fit of the Post-Thalamotomy choices in the Dundee patients (loo-log like = −0.6) than the winning Pre­Thalamotomy model (Bayes-SMEP, loo-log like = −0.68). This suggests that when the cortico­thalamic circuit is disrupted, human behaviour can be more closely aligned with, and described by, relatively simple models of reinforcement learning (RL). In contrast, in the intact thalamo-cortical system, these models only partially capture the multiple, nuanced, and complementary behavioural strategies, which humans express in their choices when making decisions in tasks such as the restless bandit. DE is one example of a behavioural strategy which can be expressed in behaviour in the intact thalamo-cortical circuit. The suppression of this type of choice, in combination with the observed improvement in model fits, points towards a generalisable role for the intact thalamo-cortical circuit, in actively *enriching the repertoire of behavioural solutions* to dynamically changing environments^66^. When disconnected, in this case, by the disruptive effects of ultrasound induced oedema, the range of behavioural options become narrowed to the extent that standard RL models, (known for their limitations in modelling flexible behaviour), ^67^ become closer representations of the algorithms that drive human decision making.

Our neuroimaging analysis of the effect of thalamotomy oedema on stay choices would support that the medio-dorsal nucleus is the most likely structure mediating this behavioural effect in our patients. We found that individual variation in the effect of thalamotomy on the proportion of stay choices correlated, with; i) the pattern of oedema into this nucleus, ii) the functional connectivity of this nucleus with the rest of the brain, iii) white matter tract connectivity between the MD nucleus and PFC. This was not the case for any other thalamic nucleus, including the clinical target (VIM) for tremor control using MRgFUS. The MD nucleus is known to have the largest reciprocal connectivity profile with the prefrontal cortex^29,32,43^. It is also increasingly recognised as a regulator of information coding within the pre-frontal cortex relevant to decision making^32,41,43^. In animal models, optogenetic activation of MD nucleus neurons promotes transition in PFC population coding which leads to switching and exploratory behaviour^68^. In humans, lesions of the MD nucleus lead to diminished behavioural flexibility and difficulty with tasks that require rapid cognitive switching which parallel those seen in our patients post thalamotomy^69^. Probabilistic analysis of the day one lesion oedema at the time of testing, demonstrated this most likely was mediated by oedema extending into an infero­lateral portion of the nucleus. The lateral, parvocellular (MDpc) portion projects to the lateral regions of the pre-frontal cortex including Brodmann area 46/9 (DLPFC, premotor and SMA/dACC)^29,30,36,70^. These are areas generally considered part of a cognitive control network which encode in neural activity predicting decisions to stay or switch ^71–73^. Cortical topography of our functional connectivity analysis showed strong positive correlations with this network whilst negative correlations were observed with the ventral, medial and orbital PFC. Interestingly, these regions project to and receive input from the medial or magnellocellar portion of the MD (MDmc)^36,37^. Lesions of the MDmc lead to premature switching and reward insensitivity ^42^, which is the opposite behavioural effect to what we observed in our patients. These contrasting behavioural effects may suggest a marked difference in the functional role of MDmc and MDpc on value-based decision making which is mediated by their distinct connectivity profiles to different regions of the PFC.

### Clinical implications

MRgFUS lesion formation has been recognised for its unpredictability. One series reported 18% incidence of lateral tail extension of the lesion into the internal capsule^74^. These are typically associated with a high rate of motor complications which are more obvious clinically than cognitive changes that might be expected from medial extension into the MD nucleus. Previous reports may therefore have under reported the impact of MRgFUS lesions on cognition by relying upon cognitive batteries insensitive to set-shifting errors analogous to the transient effects of focused ultrasound heating on the MD nucleus seen here. This is likely to be particularly relevant to patients undergoing staged bilateral lesioning where the capacity for cognitive reserve may be less robust^75,76^.

In our previous study (Gilmour et., al., (2024)^48^) when performing the restless bandit task, Parkinson’s disease patients who are apathetic are impaired at both choosing the highest valued bandit and persisting in this once it has been identified. This reward insensitivity is manifest behaviourally by an increased exploration and a lower proportion of stay choices^48^. This decision-making signature of apathy in PD is opposite to the effect of thalamotomy, and specifically oedema extension into the MD nucleus, which drove up stay choices, increased reward sensitivity, and shifted the explore-exploit trade-off towards exploitation. This raises the possibility that neuromodulation, targeting the inferolateral MD nucleus may be a viable future treatment to restore value-based learning and motivation in Parkinson’s disease. In support of this, Gilmour et., al., (2024) also identified a correlate of preserved motivation in non-apathetic PD that was suppression of thalamic activity during exploit choices. Deafferentation of thalamic negative feedback to PFC, though compensatory mechanisms, or neuromodulation, may promote consolidation of value-estimates so that the value of learnt experience can be expressed into motivated goal-directed behaviour.

### Study Limitations

Thalamotomy performed using MRgFUS deliberately targets the motor VIM nucleus. Our study relies upon the indirect extension of vasogenic oedema outside of this target and into surrounding structures, including MD nucleus which we propose mediates the behavioural effects on our task. Although unlikely, given we found no additional predictive value of including combinations of surrounding nuclei along with the MD in voxel-based analysis, we cannot definitively exclude the possibility that oedema extending to multiple nuclei led to the observed change in task behaviour. Future studies should aim to address this by selective, anatomically circumscribed, focused neuromodulation of the MD nucleus to replicate the behaviour seen here.

We also observed lower levels of directed exploration at baseline in the ET patients compared to healthy controls. The reasons for this disease effect are unclear. A single study reported deficits in the WCST in ET patients^77^ which was interpreted as evidence of subclinical frontal lobe dysfunction.

However, all patients studied here were screened for cognitive impairment pre-operatively making the overall conclusions of our study likely to be generalisable to the healthy brain. Replication studies of the effect observed here would benefit from con-current neuropsychology assessment to delineate this relationship further.

## Conclusions

By prospectively studying patients undergoing ultrasound thalamotomy, we demonstrate a causal role for the thalamus in shaping the cognitive strategy required for flexible, reward-based learning.

Following thalamotomy, increased reward sensitivity was accompanied by reduced choice flexibility, suggesting that the thalamus—particularly the mediodorsal (MD) nucleus—can modulate the degree to which prefrontal value models influence decision-making. This, in turn, supports the idea that an intact thalamus biases behaviour away from over-reliance on such value representations. Under conditions of environmental volatility and decision uncertainty, this thalamic function may help calibrate decision flexibility to maximise reward acquisition^78,79^.

## Methods

### Patients and controls

Thirty-eight patients with a clinical diagnosis of Essential Tremor (7 female), age 73.9±6.4), (Table 1) were recruited from MRgFUS Thalamotomy treatment centres in Dundee (n=26) and Liverpool (n=12). One patient was excluded from further analysis as they were unable to respond within the trial response window in >25% of trials in both pre- and post-operative task sessions.

Thirty-two age and sex-matched healthy controls (17 female, age 72±3.5) were recruited using the SHARE health informatics register (https://www.registerforshare.org/). Healthy controls were screened for a history of significant neurological or psychiatric conditions. Nine of the healthy controls had been tested as part of a previously published study to test a different hypothesis^48^.

The study was approved by the local ethics committee (North East Scotland 21/ES/0035). Written consent was obtained from all patients and controls in accordance with the declaration of Helsinki.

### Procedure

Patients performed the restless bandit task on a laptop twice. In all patients the first session was performed within the 24 hours before their thalamotomy. Due to differences in the clinical workflow between the MRgFUS sites (inpatient treatment in Dundee, day-case in Liverpool) Dundee patients performed the post-operative task 24 hours after their procedure. Liverpool patients were tested 2-3 hours after their thalamotomy. Healthy controls performed the task once.

### Experimental design

Patients and controls performed the restless four-armed bandit task^48^. Subjects were given written instructions on how to perform the task and were informed that with each trial they could win between 0 and 100 points and agreed to maximise outcome points.

Each trial started with presentation of four different coloured squares with all four bandits levers in the upright position representing the four choice options (Fig.1). Subjects made their selection using a four key mini-keyboard (Ecarke-EU) with the colour of each button corresponding to a square presented on the computer monitor corresponding to each of the “bandits”. The colour of each bandit (red, green, blue and yellow) and its position on the screen remained fixed between, and across, testing sessions. If a button press was not made within a 1.5 second response deadline, a large red “X” was displayed for 4.2 seconds at the centre of the screen. These trials were designated as missed trials and no outcome feedback was provided. For choices made within the response deadline, the chosen bandit was highlighted by displaying it’s lever as depressed, and a checkerboard pattern within the bandit’s centre. After a 3 second waiting time, the checkerboard pattern was replaced by the number of points earned on that trial, which were displayed for 1 second. Then the bandit image disappeared and was replaced by a fixation cross until 6 s after the trial onset, followed by a jittered inter-trial interval (Poisson distribution with mean of 2 s (0–5 s)) before the next trial was started. The payout (outcome) schedule of each of the four bandit choices varied according to a decaying Gaussian random walk. We used three instantiations from Daw et. al., (2006) ^14^ for the two experimental sessions. To prevent unlikely chance of between session learning effect, we randomly assigned different walk to each patients testing session using the *randi* function in Matlab (whereas healthy controls performed one task).

The task consisted of 300 trials. Patients and controls were supervised and given short breaks (3 minutes) after every 75 trials to improve concentration and task engagement.

### Analysis of behavioural performance

Model free metrics of behavioural performance in the task included best bandit choice (i.e. choosing the option with the most points), decision time and the probability of missing a trial. A choice made to the same bandit on two consecutive trials was defined as a stay choice. Decision time was defined as the time between the four bandits being presented and the subjects button press on that trial. Where a measure of behavioural performance is expressed as a probability, this was achieved by dividing by the total number of trials correctly executed, either within a 50-trial block, or across the whole task.

### Statistical analysis

We used a linear mixed-effects (lme) model using MATLAB *fitlme* function to examine the effects of the thalamotomy on behavioural performance across repeated task blocks. The dependant variable was behaviour (e.g. stay probability, best bandit choice, reaction time), measured across six, 50-trial blocks per subject. Group (patients pre- and post-thalamotomy, plus or minus healthy controls) were included as a fixed effect in all models. To account for the repeated measures within patients and controls, a Subject variable, *i*, was included as a random effect with both a random intercept *(b_oi_)* and a random slope *b*_1*i*_ for the effect of block 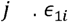 is the residual error.

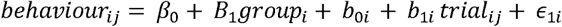

Where the influence of timing of the behavioural data collection differed (Liverpool and Dundee) an additional fixed effect of site was included.

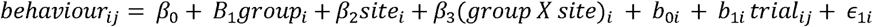

Significance of fixed effects were evaluated using F-tests with Satterthwaite’s approximation for degrees of freedom. Post-hoc effects were evaluated using linear contrasts of the fixed effects coefficients using MATLAB *coefTest* function.

All results are reported as mean values ± S.E.M. Bayesian distributions are expressed as mean posterior, prior and posterior difference of the mean with 95% higher density intervals. Correction for multiple comparisons was performed using the Bonferroni method.

A post hoc power analysis based on the Wald F test for the group × site interaction in the linear mixed-effects model indicated an estimated power of approximately 80% to detect the observed effect (1 - β ≈ 0.8 at α = 0.05).

### Computational modelling of decision making

To determine whether the thalamotomy modified decision making based upon neural value estimates of the bandits, we fitted eight computational model variations of the decision process^9^.

Each model variant used one of two learning rules, (Delta rule, or Bayesian learner) combined with one of four choice rules: i) Softmax (SM), ii) Softmax with exploration bonus (SME), iii) Softmax with perseveration bonus (SMP), (iv) Softmax with exploration and perseveration bonuses (SMEP). Posterior parameter distributions were estimated for each subject for each of the free parameters specific to the learning *(a* in the Delta learning rule) and choice rules for each model variant *(β*,φ, *p)*.

The probability Pof choosing a particular bandit, *i*, on trial, t, was determined by passing the estimated payout value, 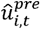 through a SoftMax function. This process was governed by four different choice rules.

*Choice rule 1* (SM):

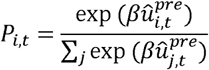

utilises a temperature parameter, *β,* to weigh choices based on their estimated value, balancing exploitation and exploration. The inverse temperature parameter, *β*, weights the sensitivity of the choices by their estimated value. Accordingly, this parameter proportionally scales the degree of exploitation versus exploration, as high values of *β* drives a greedy choice policy that favours bandits with the highest expected value 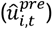.

*Choice rule 2(SME):-*

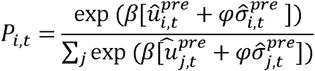

includes an additional term to further weight choices by the estimated precision (variance) of each bandit’s value, 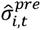. This is done by using an exploration bonus, φ, which augments uncertainty driven directed exploration.

*Choice rule 3 (SMP):-*

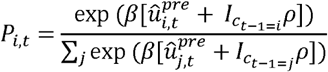

Includes a term to weight each bandit’s choice probability in the model by a perseveration bonus, *p* ^16,17^, which is added to the chosen actions probability of being chosen, if two more consecutive trials are repeated, *I_ct-_*_1=*i*._

*Choice rule 4 (SMEP):-*

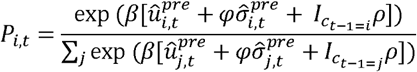

A fourth choice rule included both the exploration and perseveration bonuses.

In both the SMP and SMEP choice rule (rules 3 & 4), the variable represented by *I*, is either 0 or 1 depending upon whether the chosen bandit on trial *t* is the same as that on the previous trial *c_t___1_*.

### Bayesian Learner

The Bayesian learner implements the Kalman filter ^18^ to keep track of the estimated value 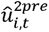 and uncertainty 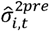 of each bandit *i* on trial *t*. We initialise these beliefs as 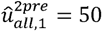 and 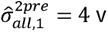 ^13^. After a bandit is chosen *c_t_* the reward for that bandit *i* is obtained *r_t_.* The prediction error *δ_t_* is

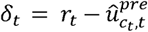

The Kalman filter, has a variable learning rate (the Kalman gain), where the confidence in the measurement makes a dramatic difference to the amount learned from it. The estimated variance of the given action 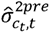 and the observation variance 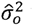 which represents the uncertainty in all observations. 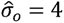 for the purpose of fitting due to model degeneration if freed, a problem previously encountered _1_^3,14^. The Kalman gain *κ_t_* is given as:

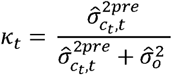

The Kalman gain naturally increases, such that options that have high uncertainty can be quickly estimated, but options that are well established are fine-tuned, with only a small amount learned from trial to trial. The estimated value of the bandit 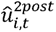 and the associated variance 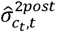 can then be calculated as:

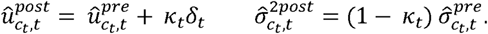

For bandits that are not chosen on any one trial, the prior mean and variance remains unchanged within a trial. However, the prior distributions are updated for all bandits *in between* trials based upon the subject’s belief about the gaussian random walk such that;

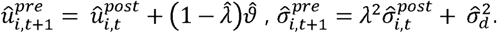

Where 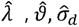 are constants representing the decay parameter, decay centre and the diffusion variances and were fixed at values of 0.98, 50, and 2.8 for each respective parameter. We used the same values as Chakroun et. al, (2020^13^) which are derived from the actual parameters that govern the gaussian walk underlying the bandit’s payout schedule.

### Delta rule

The delta learning rule has a fixed learning rate, *a*,and there is no variance tracking for the uncertainty of the valuation. After a bandit is chosen *c_t_* the reward for that bandit *i* is obtained *r_t_.* The prediction error *δ_t_* is derived in the same way and the prediction of the bandit’s value 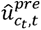 can be updated as such:

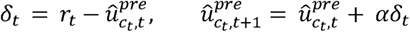

There is no decay in this model between trials, the estimated value for a bandit is only updated when that bandit is selected. In the absence of an equivalent estimate of uncertainty 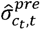 in the delta learner, we modified choice rules 2 and 4 (SME, SMEP) so that the exploration bonus *φ* was modified by the how long ago a bandit *i* was chosen *t_i_*;

*Choice rule 2(SME):*

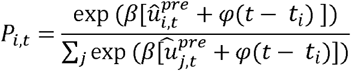

*Choice rule 4 (SMEP):*

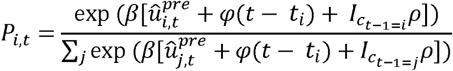

### Model Fitting

Posterior parameter distributions were estimated for each subject for each of the free parameters specific to the learning *(a* in the Delta learning rule) and choice rules for each model variant *(β,φ, p)*.

Parameter estimates were derived using hierarchical Bayesian modelling using Stan within CmdStan (v2.36.0) via CmdStanPy (v1.2.5) in Python 3.13.5. Sampling was performed with four chains, each chain running after a warmup period of 2500 iterations for a total of 10000 samples. The prior for each group-level mean was uniformly distributed and bounded. For each group-level standard deviation, a half-Cauchy distribution with location parameter 0 and scale parameter 1 was used as a weakly informative prior ^19^. Priors for all subject-level parameters were normally distributed with a parameter­specific mean and standard deviation.

### Lesion mapping analysis

We acquired both pre- and post-operative imaging using a 3T Siemens Prisma Fit Scanner (Siemens Healthineers, Erlangen, Germany) and proprietary 20 channel head and neck coil. Data used in lesion mapping analysis included: Pre-operative volumetric T_1_-weighted images using a MPRAGE sequence; Diffusion tensor imaging (DTI/dMRI) acquired pre-operatively using a spin echo – EPI sequence. Post-operative demarcation of the thalamotomy used a T_2_-weighted FLAIR-SPACE sequence. Magnetization Prepared Rapid Gradient Echo (MPRAGE) sequence, [Echo Time (TE)/repetition time (TR) = 2.32/2300 ms, inversion time (TI) = 900 ms, flip angle = 8°, field of view (FOV) = 240 mm, isotropic voxels 0.9x0.9x0.9mm3, acquisition time (TA) = 5min21sec]. Diffusion tensor imaging (DTI/dMRI) acquired pre- and post-operatively using a spin echo, echo planar imaging sequence (EPI), [diffusion weighting b = 0 s/mm^2^ and in 30 uniformly distributed directions (b = 1000 s/mm^2^) and TE/TR = 93.0/3100ms, averages = 4, FOV = 230 mm, voxels = 1.8x1.8x5.0mm3, TA = 6min35sec]. Post-operative T_2_-weighted Fluid attenuated inversion recovery Sampling Perfection with Application optimized Contrasts using different flip-angle Evolutions (FLAIR-SPACE) sequence [TE/ TR = 387 /5000 ms, TI = 1600 ms, FOV = 230 mm, voxel size = 0.4 x 0.4 x 0.9 mm3, TA = 5min 42sec].

Patients treated in Liverpool where MRI was performed within hours of their procedure were not included in the lesion-behaviour mapping as full maturation of the thalamic oedema does not occur and is not maximal until at least 24hours post proceedure^47^. We therefore limited our analysis mapping the relationship of this factor and changes in task behaviour, to the patients treated in

Dundee, for whom post-operative MRI and behavioural data were acquired concurrently, allowing oedema definition on MRI. One patient’s post operative imaging in the Dundee group was not available due to technical issues with the MRI scanner.

Post-operative T_2_-weighted images were used to manually delineate the voxels enveloped by thalamotomy oedema to create a binary mask of the lesion. The thalamotomy mask was demarcated to include the necrotic lesion core (hypointense), surrounding cytotoxic oedema (markedly hyperintense) and the outer margin by the surrounding perilesional oedema (moderately hyperintense)^80^. Masking was performed following normalisation into Montreal Neurological Institute (MNI) space using SPM12 (https://www.fil.ion.ucl.ac.uk/spm/software/spm12).

Lesion masks were subject to three further post-processing analyses: i) probabilistic voxel-wise statistical mapping of oedema overlap with individually segmented motor and higher-order thalamic nuclei^28,81–84^, ii) functional connectivity analysis using normative resting-state group connectivity^85–87^, iii) white matter connectivity from individual preoperative DTI, reconstructed using probabilistic tractography^88–92^.

### Probabilistic voxel-wise statistical mapping

Following manual delineation of the thalamotomy oedema margins a “lesion-nucleus overlap” mask was generated for each patient which consisted of voxels with shared MNI co-ordinates with individual thalamic nuclei. Individual patient segmentations of the thalamus were performed using HIPS-THOMAS (Version running on Python v3. 9.13) applied to the spatially normalised pre-operative volumetric T_1_-weighted images. In addition to the targeted VIM nucleus, overlap masks were created for the MD, CM and VLa as all three have previously associated with cognitive function and are in close proximity to the VIM.

The individual patient masks for each nucleus where then subjected to voxel-wise statistical mapping to define probability mask by averaging across voxels in MNI space. To define the “sweetspot” of maximal behavioural influence of thalamotomy, we weighted each patient’s mask with the measured behavioural outcome and subjected each voxel to a Wilcoxon signed rank test. Correction for false discovery rate used a *q* value of 0.05 applying the Benjamini & Hochberg method for multiple comparisons.

Using MATLAB’s *glmfit*, a GLM was applied to each voxel with the change in the patients P(Stay) (pre-post thalamotomy) as the response and the binary mask (1 = oedema present, 0 = no oedema) the predictor variable. To avoid edge effects of variation in the individual thalamic nuclear anatomy between subjects, we limited our analysis to voxels where at least≥25% of the patients had overlap between their individually segmented nuclei. Voxels were considered significant where their Beta weights equalled or exceed the 95% confidence limits of a two-tailed permutation distribution. To correct for multiple comparison’s we identified spatially contiguous, statistically significant voxels using connected component labelling (CCL). The mean Beta weight of voxels forming a cluster was then tested for cluster-level significance by comparing their values against a two-tailed permutation distribution (permutation per cluster n = 5000). For both the voxel and cluster-based permutation the permuted estimates were derived by re-estimating the GLM Beta weights after randomly shuffling the order of each patients change in P(Stay). The number of permutations per voxel and per cluster was 1000 and 5000 respectively. Clusters whose mean Beta weights could not be explained at p(perm)<0.05 were considered statistically significant.

### Functional connectivity behavioural analysis

Using the THOMAS thalamic atlas in Lead-DBS V2.5 (https://www.lead-dbs.org), each patient’s lesion oedema-nucleus overlap mask was used as seed regions to estimate individual functional connectivity profile maps. Functional connectivity was estimated using the openly available group connectome derived from resting state functional connectivity MRI data of 1000 healthy volunteers GSP1000 connectome (Holmes et. al., 2015) . Lead DBS includes pre-compiled averaged connectivity matrix estimated from 1000 rs-fMRI which is a voxel-by-voxel connectivity values of every voxel combination within MNI space. Using the thalamotomy overlap-masks as the seed region, the functional connectivity is calculated by summing the connectivity values from all voxels within this seed region to each voxel with the normative connectivity matrix. We then used each patient’s connectivity map to perform voxel-wise correlations between the estimated connectivity profile and the change in behaviour in the restless bandit task. The resulting “R-map” therefore represents the visualisation of brain regions where high levels of functional connectivity correlated (either positively or negatively) with the behavioural effect of thalamotomy. To test the validity of this map’s predictions, we correlated each individual’s connectivity profile with the R-map’s correlation coefficients across every brain voxel. This produced an estimate of the spatial similarity between the patient’s functional connectivity profile and the group level R-map. For the predictive value of an R-map to be meaningful, the change in behaviour should correlate linearly with the spatial similarity of the individuals connectivity to the original R-map. To test for correlations that might arise by chance, we calculated spatial similarly estimates after 1000 permutations of the behavioural data and re-estimated this the correlation coefficient. This analysis was repeated for each thalamic nucleus (MD, VIM, VLa & CM). This allowed us to build a profile of how much individual variation in the effect of thalamotomy on behaviour, could be explained by the functional connectivity profile of oedema extension into each nuclei.

### Probabilistic tractography analysis

Patients pre-operative T_1_-weighted and post-operative T_2_-weighted images were co-registered using lead DBS. The co-registered pre-operative MRI was then used to segment pre-frontal cortical regions and individual thalamic nuclei using Freesurfer. Transforming the same binary masks used in the functional connectivity analysis, back into native space, we then created a second mask, ‘MD-thalamotomy overlap’, which consisted of voxels where thalamotomy oedema overlapped with the patients MD nucleus.

Pre-operative dMRI images were subject to skull stripping, bias field correction and corrected for geometric distortion using the Brainsuite Diffusion Pipeline(https://brainsuite.org/processing/diffusion). Co-registration of the dMRI to the pre-operative T_1_-weighted image, and then registration to MNI space were performed using Advanced Normalisation Tools. White matter fibre orientations were estimated using FSL’s *bedpostx* and streamline densities using *probtrackx*. For each voxel we generated 5000 streamlines, with a step-length of 0.5mm a curvature threshold of 0.2, a discard threshold of 0mm and 2000 number of steps per sample. To control for number of voxel differences, we normalised the number of streamlines by the number of voxels in the seed mask.

The MD-thalamotomy overlap mask and a Freesurfer segmented ROI within the prefrontal cortex (frontal pole and rostral middle frontal gyrus) were defined as the seed and waypoint regions, respectively, for probabilistic tractography. Cortical ROI’s were defined using the Desikan-Killiany Atlas. Individual streamline density map’s were wrapped back into MNI space before voxel-wise GLM analysis with FSL’s *randomise* function using a one-sample T-Test or a regression design matrix. Clusters were defined using Threshold-Free Cluster Enhancement (TFCE) after 5000 permutations.

## Data Availability

Behavioural data from these experiments and python scripts used for computational modelling are openly available at https://osf.io/2wzqy/

## Acknowledgements

The authors would like to thank both the DBS team at the Queen Elizabeth University Hospital, Glasgow (Dr’s Vicky Marshall, Michael Canty & Edward Newman) and the MRgFUS team of the Walton Centre, Liverpool (Dr Jay Panicker, Dr Dinesh Damodaran, Dr Mark Radon & Dr Rajesha Srinivasaiah) who’s collective clinical efforts in patient selection allowed this research to be undertaken. We would also like to thank the patients who participated in this study.

