## Supplementary Information for "Focused ultrasound neuromodulation of mediodorsal thalamus disrupts decision flexibility during reward learning"

**Supplementary Results**

*Computational modelling – model comparison*

| Loo log-lik | AlphaSM | AlphaSME | AlphaSMP | AlphaSMEP | BayesSM | BayesSME | BayesSMP | BayesSMEP |
| --- | --- | --- | --- | --- | --- | --- | --- | --- |
| all | -0.783 | -0.766 | -0.664 | -0.653 | -0.770 | -0.742 | -0.660 | **-0.647** |
| Pre Thal | -0.815 | -0.797 | -0.696 | -0.691 | -0.800 | -0.772 | -0.692 | **-0.687** |
| Post Thal liverpool | -0.821 | -0.799 | -0.680 | -0.679 | -0.792 | -0.774 | -0.681 | **-0.663** |
| Post Thal  Dundee | -0.630 | -0.605 | -0.474 | -0.467 | -0.615 | -0.571 | -0.470 | **-0.458** |
| HC | -0.850 | -0.841 | -0.767 | -0.743 | -0.846 | -0.826 | -0.761 | **-0.740** |

| BIC | AlphaSM | AlphaSME | AlphaSMP | AlphaSMEP | BayesSM | BayesSME | BayesSMP | BayesSMEP |
| --- | --- | --- | --- | --- | --- | --- | --- | --- |
| all | 0.452 | 0.457 | 0.561 | 0.566 | 0.461 | 0.471 | 0.570 | **0.583** |
| PreThal | 0.438 | 0.443 | 0.545 | 0.546 | 0.447 | 0.457 | 0.555 | **0.563** |
| PostThal  Liverpool | 0.434 | 0.442 | 0.547 | 0.547 | 0.450 | 0.457 | 0.558 | **0.570** |
| Post Thal  Dundee | 0.527 | 0.537 | 0.662 | 0.664 | 0.538 | 0.559 | 0.673 | **0.684** |
| HC | 0.423 | 0.424 | 0.514 | 0.528 | 0.427 | 0.433 | 0.522 | **0.541** |

| MLT | AlphaSM | AlphaSME | AlphaSMP | AlphaSMEP | BayesSM | BayesSME | BayesSMP | BayesSMEP |
| --- | --- | --- | --- | --- | --- | --- | --- | --- |
| all | -0.779 | -0.760 | -0.607 | -0.592 | -0.767 | -0.737 | -0.602 | **-0.578** |
| PreThal | -0.810 | -0.792 | -0.637 | -0.628 | -0.797 | -0.768 | -0.632 | **-0.616** |
| PostThal  Liverpool | -0.818 | -0.792 | -0.626 | -0.619 | -0.790 | -0.768 | -0.621 | **-0.595** |
| Post Thal  Dundee | -0.624 | -0.598 | -0.428 | -0.417 | -0.612 | -0.566 | -0.423 | **-0.403** |
| HC | -0.845 | -0.834 | -0.704 | -0.674 | -0.843 | -0.821 | -0.697 | **-0.664** |

**Supplementary Table 1 Model Comparison** *Loo-log-lik; Leave-one-out cross validated Log likelihood; BIC; Bayes Information Criteria; MLT; Mean Likelihood per trial. SM; Softmaxl SME; Softmax with Exploration bonus; SMEP; Softmax with Exploration and perseveration bonuses. Values in bold are the winning model for each group.

|  | **Model Parameter** | | | | | | | | |
| --- | --- | --- | --- | --- | --- | --- | --- | --- | --- |
| **Group**  **Posteriors** | $\beta$ (reward sensitivity) | | | $\varphi$ (exploration bonus) | | | $\rho$ (perseveration bonus) | | |
|  | **Mean** | **95% HDI** | | **Mean** | **95% HDI** | | **Mean** | **95% HDI** | |
| **Pre-Thalamotomy** | 0.104 | 0.098 | 0.109 | 0.463 | 0.238 | 0.667 | 11.213 | 11.205 | 12.181 |
| **Post-Thalamotomy**  **Liverpool** | 0.118 | 0.107 | 0.131 | 0.888 | 0.556 | 1.212 | 11.834 | 10.241 | 13.365 |
| **Post-Thalamotomy**  **Dundee** | 0.144 | 0.133 | 0.156 | 0.627 | 0.408 | 0.819 | 12.345 | 11.311 | 13.419 |
| **Healthy Control** | 0.105 | 0.099 | 0.111 | 0.870 | 0.638 | 1.073 | 10.559 | 9.417 | 11.613 |
| **Contrasts**  **(Post Dundee minus Pre)** | **0.040** | **0.030** | **0.050** | 0.164 | -0.137 | 0.470 | 1.136 | -0.263 | 2.26 |
| **Contrast**  **(Post-Dundee**  **Minus**  **Post-liverpool)** | **0.025** | **0.018** | **0.042** | -0.261 | -0.542 | 0.043 | 0.51 | -1.368 | 2.404 |
| **Contrast**  **Post-Liverpool**  **Minus**  **Pre)** | **0.015** | **0.002** | **0.028** | **0.425** | **0.0255** | **0.797** | 0.620 | -1.246 | 2.464 |
| **Contrast**  **HC - pre** | -0.001 | -0.01 | 0.006 | **-0.406** | **-0.720** | **-0.106** | 0.65 | -0.805 | 2.147 |

**Supplementary Table 2 Summary of Posterior distributions and contrast estimates for parameters of the wining model for each group**. *Denotes Significant differences in the posterior distributions of the parameter estimate at 95% confidence limit.

*Neuroimaging*

|  | **Whole Thalamotomy** | **MD** | **Vim** | **CM** | **VLa** |
| --- | --- | --- | --- | --- | --- |
| **Number of Voxels** | 2102±241 | 162±21 | 229±16 | 83±4 | 36±7 |
| **Correlation**  **Change in P(Stay)** | -0.32  p=0.1 | -0.05  p=0.81 | -0.08  p=0.70 | -0.25  p=0.23 | 0.26  p=0.21 |
| **Correlation**  **Change in**  **P(Exploit)** | -0.11  p=0.58 | -0.06  P=0.76 | -0.07  p=0.74 | 0.31  p=0.13 | 0.16  p=0.43 |

**Supplementary Table 3 Correlations between thalamotomy oedema extension into individual thalamic nuclei and task behaviour.** Voxel oedema values are expressed as mean±s.e.m Correlations coefficients are Pearson’s rho values with corresponding Bonferroni adjusted p-values. MD, Mediodorsal nucleus, VIM, Ventral-intermediate, CM centromedian, VLa, ventrolateral anterior.

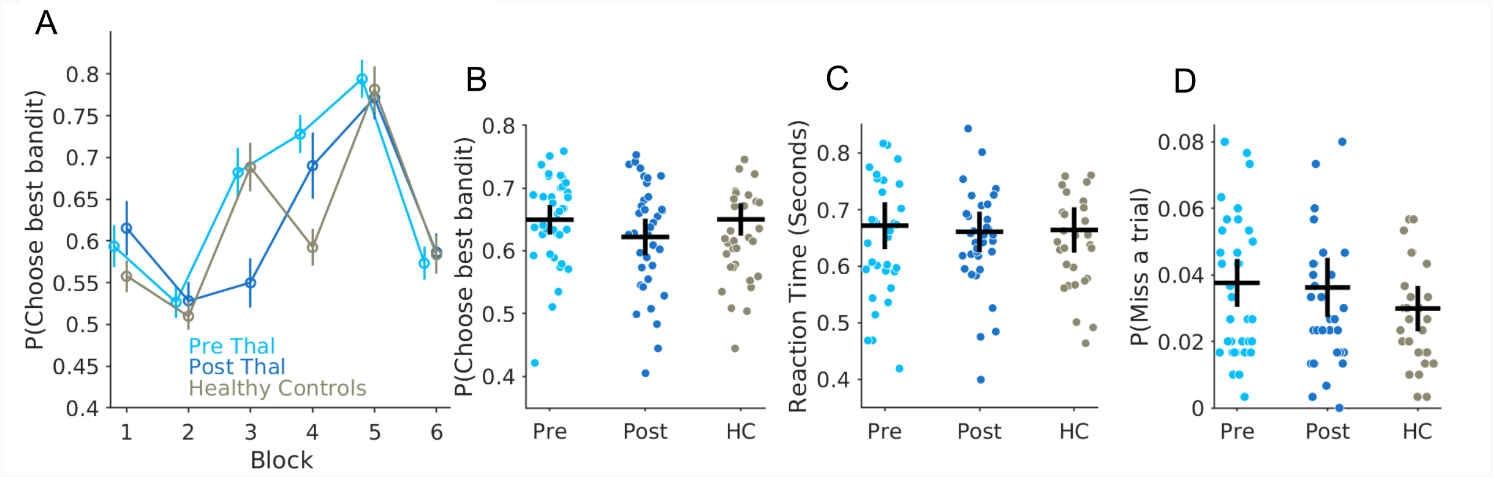

**SFig 1. Performance in the restless bandit task is unaffected by Thalamotomy.** Average probability of choosing the bandit with the highest payout, P(Choose best bandit) is plotted pre (light blue) and post-thalamotomy(dark blue) and in an healthy control group (grey). (**A**) The increase in average values of best bandit choice plotted across six, 50 trial blocks across the task. Vertical lines S.E.M. In **(B**) each circle represents average best bandit choice probability across the task for an individual subject with the horizontal and vertical bar represents the group mean and 95% confidence limits. Average reaction time (**C**) and proportion of trials where the response window was missed (D) where unaffected by the thalamotomy. No significant differences were found between the HC and pre-thalamotomy performance in the proportion of best bandit choices, reaction time or proportion of missed trials.

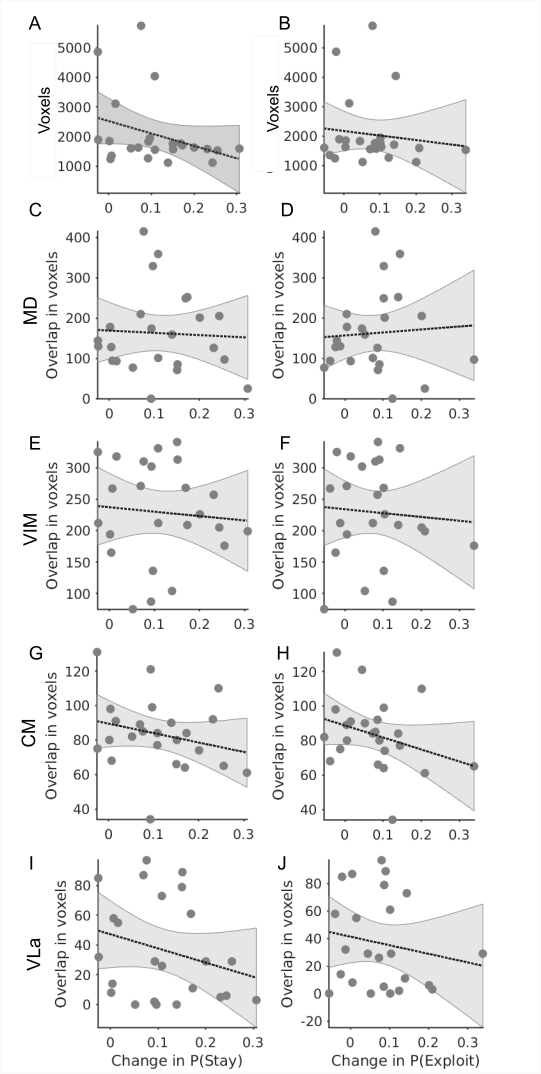

**SFig2. The total volume of thalamotomy oedema and overlap into individual thalamic nuclei do not correlate with changes indecision making behaviour.** (**A**) Correlation between the total number of oedematous voxels caused by the whole thalamotomy and the probability of making a stay choice in the task (P(Stay) and change in P(Exploit), (**B**). Correlations between change in P(Stay) over the task, pre- minus post-thalamotomy, and the number of oedematous voxels from the thalamotomy that overlap into each segmented nucleus surrounding the surgically targeted Vim (segmentation based upon Su et. al., (2019) including the MD (C), VIM (E), CM (**G**),and VLa (I). The same analysis applied to each nuclei for the change in P(Exploit) is illustrated in (**D**), (**F**), (**H**) & (J).

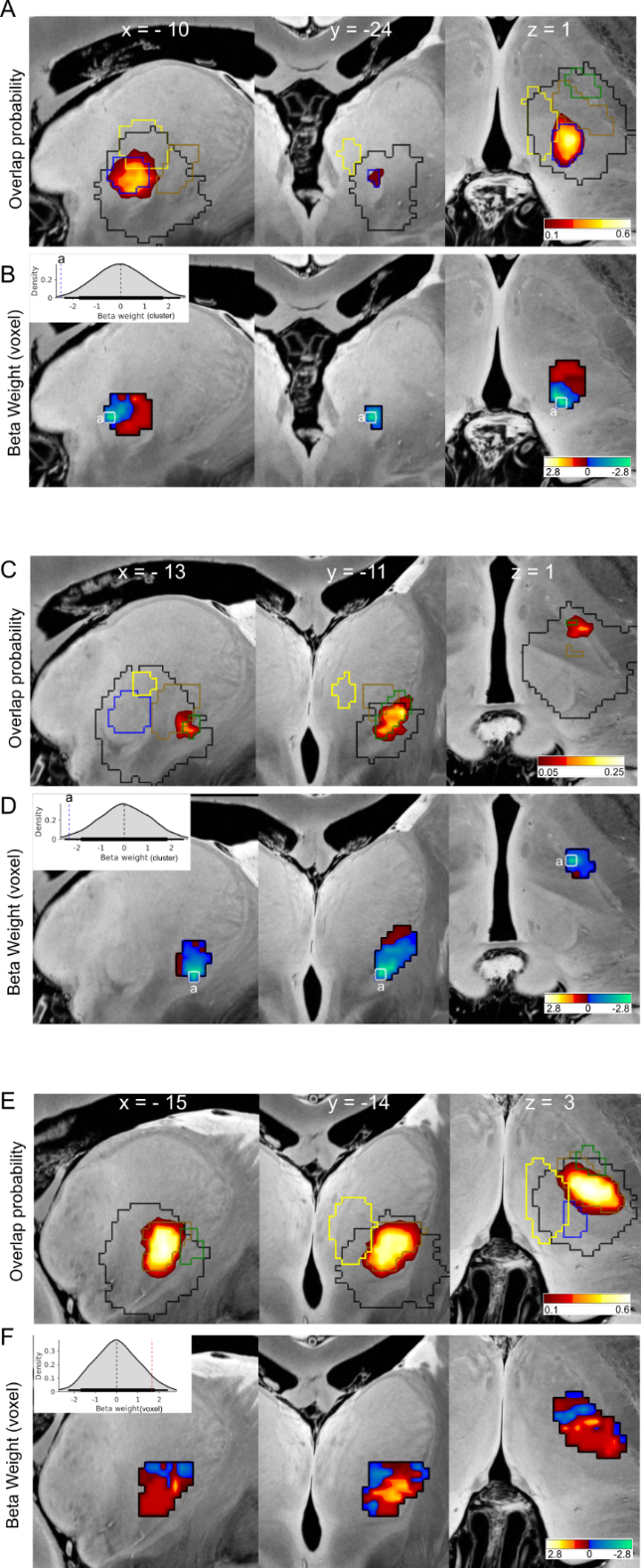

**SFig3. Voxel Wise GLM analysis of thalamotomy oedema effect on P(Stay).** Sagittal, coronal and axial slices with the yellow-red heatmap illustrate the probability of lesion oedema extending into the each nucleus. CM (A), VLa (C) and VIM (E)The outer border of the MD (yellow), CM, (blue) VIM (brown) and VLa (green) are superimposed alongside the outer oedema margins in black. The corresponding Beta weights from a GLM, fitting the presence of oedema in voxel to the individual patients change in P(Stare illustrated in (B), (D) and (F) for the each respective nuclei. The outer border of significant clusters are outlined in white and the cluster- based permutation distribution inlayed in grey. .Horizontal line represent 95% (thick) and 99% (thin) confidence limits.

| Mediodorsal Nucleus (MD) | | | | | | | |
| --- | --- | --- | --- | --- | --- | --- | --- |
| Cluster size (Voxels) | Beta weight  (voxel  peak) | Beta weight  (cluster  average) | Permutation corrected  Cluster  P-value | Permutation corrected  Voxel  P-value | Peak  MNI co-ordinate (mm) | | |
|  |  |  |  |  | X | Y | Z |
| 14 | -2.6 | -1.86 | 0.004 | 0.002 | -7 | -21 | -1 |
| 7 | -2.13 | -1.53 | 0.005 | 0.01 | -8 | -21 | 5 |
| 4 | 2.75 | 2.17 | 0.01 | 0.01 | -8 | -19 | 6 |
| Ventrolateral anterior nucleus (Vla) | | | | | | | |
| 5 | -2.39 | -2.33 | 0.02 | 0.03 | -13 | -11 | 1 |
| Ventral Intermediate nucleus (Vim) | | | | | | | |
| n/a | -1.73 | n/a | n/a | 0.13 | -9 | -13 | 0 |
| n/a | 1.66 | n/a | n/a | 0.11 | -15 | -14 | 3 |
| Centromedian (CM) nucleus | | | | | | | |
| 3 | -2.59 | -2.50 | 0.01 | 0.01 | -10 | -24 | 1 |

**Supplementary Table 4 Results of Voxel wise GLM analysis of P(Stay) on MD and surrounding nuclei.**

| *Positive Peak Coordinates X/Y/Z (Value)* | *Negative Peak Coordinates X/Y/Z (Value)* |
| --- | --- |

| DLPFC | RH | 37/45/32 (0.54) |
| --- | --- | --- |
|  | LH | -38/46/32 (0.51) |
| Pre SMA | RH | 6/9/53 (0.61) |
|  | LH | -5/10/41 (0.60) |
| Insula  (BA13) | RH | 32/-6/16 (0.62) |
|  | LH | -34/-6/-16 (0.64) |
| Premotor (BA 6) | RH | 58/4/38 (0.63) |
|  | LH | -58/4/38 (0.67) |
| Occipital | RH | 58/24/20 (0.65) |
|  | LH | 58/24/20 (0.6) |
| CBM | RH | 21/-64/-53 (0.61) |
|  | LH | -17/62/-57 (0.65) |
| Supramarginal Gyrus  (BA40) | RH | 62/-25/46 (0.59) |
|  | LH | -62/-22/24 (0.57) |

| Frontal Pole  (BA10) | LH | -14/58/16 (-0.65) |
| --- | --- | --- |
|  | RH | 6/56/16 (-0.61) |
| dACC | RH | 2/41/-6 (-0.67) |
|  | LH | -5/-41/-5(-0.66) |
| OFC (47) | LH | -37/31/ -17(-0.61) |
|  | RH | 35/26/-17(-0.58) |

**Supplementary Table 5 Peak Rmap values and voxel co-ordinates of cortical regions who’s connectivity with the Mediodorsal thalamotomy oedema mask predicts increased stay choices.** Correlations are all peak r-values with corresponding MNI co-ordinates. DLPFC; Dorsolateral prefrontal cortex, SMA, supplementary motor area, CBM cerebellum, dACC, dorsal anterior cinguiate cortex, OFC, Orbitofrontal cortex.

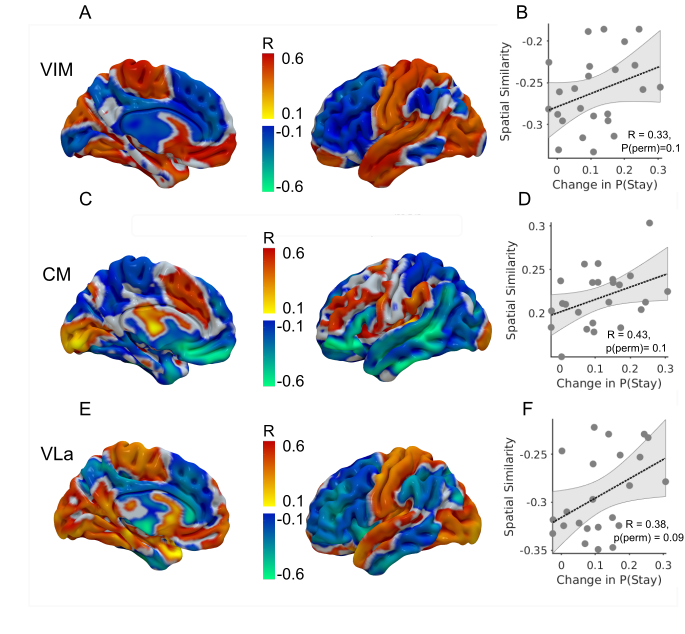

**SFig4. Rmap analysis of thalamotomy effect on P(Stay).** Medial (**A**) and lateral cortical surface projections of the whole-brain voxel-wise R- Map illustrated functional connectivity profile between thalamotomy oedema extension into the VIM nucleus and increase in Pstay). The warm and cold colour bars represent the rho value for positive and negative correlations respectively **B**) Functional connectivity “R-map” between the VIM- thalamotomy oedema did not significantly predict the individual variation in increased P(Stay). The same analysis applied to the CM (**C&D**) and VLa (**E&F**) was also unable to significantly predict this behaviour after correction for multiple comparisons following permutation testing.

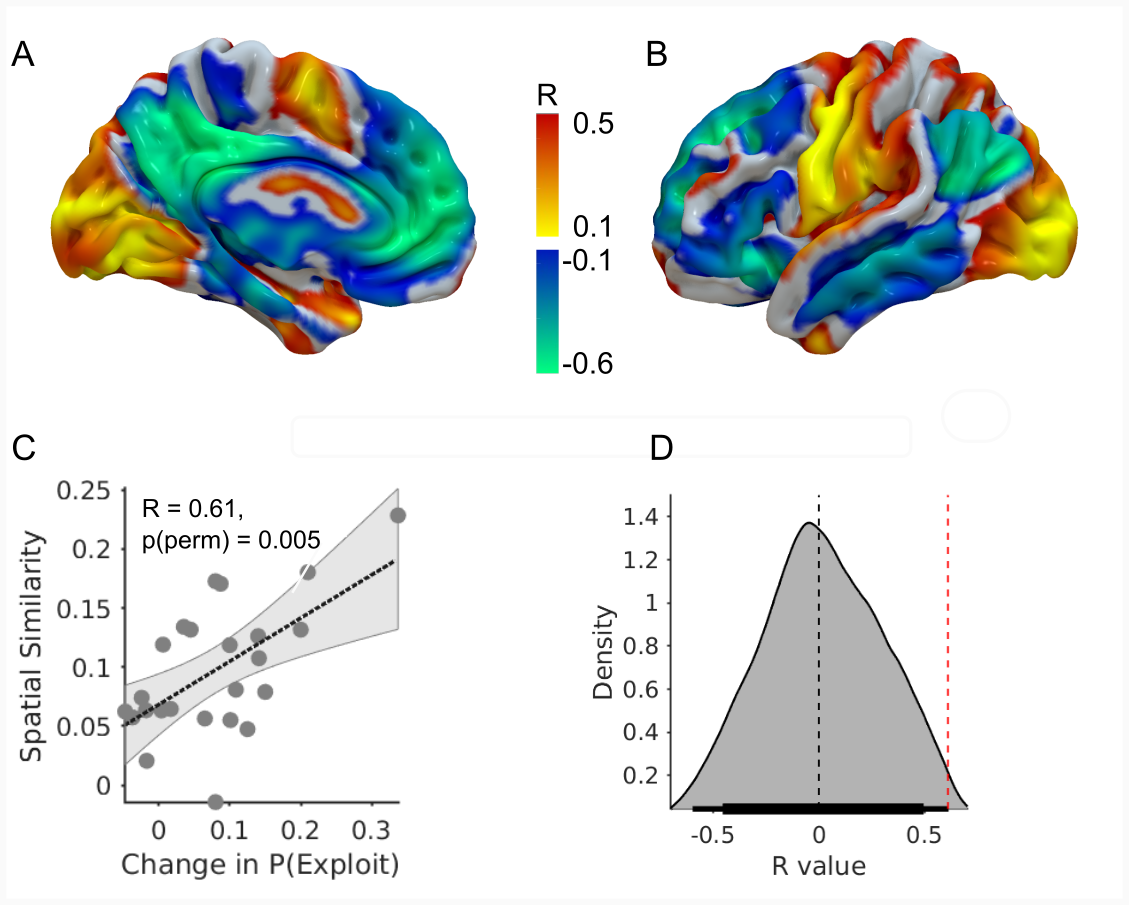

**SFig5. Mediodorsal thalamus-prefrontal cortical connectivity predicts thalamotomy effect on exploitation.** Medial (**A**) and lateral (**B)** cortical surface projections of the whole-brain voxel-wise R- Map demonstrates the optimal functional connectivity profile between thalamotomy oedema extension into the MD nucleus and increase in P(Exploit). Warm colours show cortical voxels where functional connectivity to the voxels of the MD nucleus associated with increased P(Exploit). Cool colours indicate voxels where stronger functional connectivity was associated with less marked effect of thalamotomy on the same behaviour. **(C**) The more the individual functional connectivity profile matched the ‘optimal’ R-Map, the greater was the increase in P(Exploit) induced by thalamotomy related oedema into the MD nucleus (R = 0.61, p=0.0051). In (**D**) we plot the R value distribution derived from 1000 re-permuted correlations between the individual change in P(Exploit) caused by the thalamotomy and the optimal R-map. The probability of seeing the same rrelation by chance was p<0.01 (represented by the red dashed vertical line). Horizontal thin and thick lines represent 99 and 95% HDI.(**D**).

| *Positive Peak Coordinates X/Y/Z (Value)* | *Negative Peak Coordinates X/Y/Z (Value)* |
| --- | --- |

| Premotor | RH | 53/0/52 (0.56) |
| --- | --- | --- |
|  | LH | -52/-1/45 (0.59) |
| Premotor | RH | 60/1/27 (0.58) |
|  | LH | -60/1/27 (0.61) |
| CBM | RH | 25/-58/-60 (0.55) |
|  | LH | -20/-58/-59 (0.55) |
| preSMA | RH | -10/1/61 (0.50) |
|  | LH | 11/4/60 (0.49) |

| dACC | RH | -1,42,-2 (-0.58) |
| --- | --- | --- |
|  | LH | 2/43/-2 (-0.58) |
| vACC | RH | -2,-21,38 (-0.61) |
|  | LH | 2/-23/36 (-0.58) |
| Frontal Pole (BA10) | LH | -10/58/13 (-0.59) |
|  | RH | 9/54/11(-0.55) |

**Supplementary Table 6 Peak Rmap values and voxel co-ordinates of cortical regions who’s connectivity with the Mediodorsal thalamotomy oedema mask predicts increased exploitation.** Correlations are all peak r-values with corresponding MNI co-ordinates. DLPFC; Dorsolateral prefrontal cortex, SMA, supplementary motor area, CBM cerebellum, dACC, dorsal anterior cinguiate cortex, vACC ventral anterior cingulate cortex.

| Cluster size (Voxels) | P-value (FWE corrected) | MNI co-ordinate (mm) | | | Region |
| --- | --- | --- | --- | --- | --- |
|  |  | X | Y | Z |  |
| 37884 | <0.001 | -16 | 57 | -10 | Left FP (BA10) |
| 27 | 0.012 | -34 | 56 | 6 | Left FP (BA10) |
| 10 | 0.031 | -20 | 49 | 35 | Left DLPFC (BA 9) |
| 10 | 0.031 | -11 | 42 | -23 | Left DLPFC (BA 9) |

**Supplementary Table 7 Clusters of significant group level structural connectivity between thalamotomy oedema into MD nucleus and the pre-frontal cortex.** MNI co-ordinates of peak cluster statistic. FP; Frontal Pole, DLPFC; Dorsolateral prefrontal cortex.

| Cluster size (Voxels) | P-value (uncorrected) | MNI co-ordinate (mm) | | | Region |
| --- | --- | --- | --- | --- | --- |
|  |  | X | Y | Z |  |
| 446 | 0.024 | -52 | 25 | 19 | Left DLPFC (BA 46) |
| 37 | 0.032 | -40 | 33 | 31 | Left DLPFC (BA 9) |
| 20 | 0.035 | -12 | 57 | -12 | Left FP (BA10) |

**Supplementary Table 8 Clusters of MD-PFC structural connectivity which positively correlated with increased P(Stay) choices..** MNI co-ordinates of peak cluster statistic. FP; Frontal Pole, DLPFC; Dorsolateral prefrontal cortex.

| Cluster size (Voxels) | P-value (uncorrected) | MNI co-ordinate (mm) | | | Region |
| --- | --- | --- | --- | --- | --- |
|  |  | X | Y | Z |  |
| 178 | 0.019 | -19 | 44 | -20 | Left OFC (BA 11) |
| 175 | 0.023 | -39 | 43 | 26 | Left FP (BA10) |
| 79 | 0.038 | -16 | -5 | -3 | Left GPI |
| 30 | 0.018 | -12 | 19 | -4 | Left Caudate |
| 19 | 0.046 | -11 | -18 | -3 | Left Midbrain |

**Supplementary Table 9 Clusters of MD-PFC structural connectivity which positively correlated with increased P(Exploit) choices.** MNI co-ordinates of peak cluster statistic. FP; Frontal Pole, OFC; Orbitofrontal cortex.GPI; Globus Pallidus Interna.

**Supplementary Information**

***Therapeutic ultrasound treatment parameters***

We report treatments in line with recommendations for reporting therapeutic ultrasound treatment parameters(Padilla & Ter Haar, 2022).

**Focused Ultrasound Transducer & System:**

MRgFUS VIM thalamotomy was performed using the Insightec Exablate Neuro 4000 (Insightec, Haifa, Israel, S/N: 4226) mounted on a Siemens 3T Magnetom Prisma (Siemens Healthineers, Erlangen, Germany). The transducer helmet is a hemispherical 1024 element array with a 30.0 cm diameter and 15.0 cm focal length, operating at a central frequency of 670 kHz.

Initial target localisation is performed preoperatively, off-console by consultant neurosurgeon (SK) using preoperative MR acquired on the Siemens 3T Magnetom Prisma to localise the VIM. These include volumetric T1 weighted Magnetisation Prepared Rapid Gradient Echo (MPRAGE) sequence (TE/TR (echo time to repetition time) of 2.32 ms / 2300 ms, inversion time (TI) of 900 ms, flip angle 8º, field of view 240 mm, isotropic voxels of 0.9 mm × 0.9 mm × 0.9 mm) and Diffusion Tensor Imaging (DTA/dMRI) (using spin echo, echo planar imaging sequence with TE/TR = 93.0/3100 ms, averages =4, FOV = 230 mm, voxels 1.8 mm × 1.8 mm × 5.0mm, diffusion weighting b = 0 s mm^-2^and in 30 uniformly distributed directions b = 1000 s mm^-2^).

On the Exablate console, treatment planning uses the WMn preoperative scan is fused with preoperative CT (GE Revolution EVO CT; BONEPLUS convolution kernel, in-plane spacing of 0.467 mm × 0.467 mm, slice thickness of 0.625 mm and slice spacing of 0.625 mm). CT is used to mask areas of calcifications and calculate skull density ratio (the ratio of the density of trabecular to cortical bone).

On the day of the operation, the patient has their head shaved and is fixed into a stereotactic head frame (Exablate 4000 Headframe, Insightec, Haifa, Israel). The patient is then taken into the MR suite and fixed relative to the MR scanner within the transducer helmet, which is mounted on the Siemens Prisma 3T. The space between the transducer and helmet is filled with degassed, cooled water (~14 ºC)

An anatomical scan is acquired (3D CISS, TR/TE 7.79/3.99, flip angle 41d, in-plane pixel spacing 1.11 mm × 1.11 mm, nominal slice thickness 1.5 mm) and this is fused with preoperative WMn and CT. The system uses the fused CT to calculate skull-correcting phase shifts per element (via proprietary algorithm). The initial target coordinates are located within this fused system, and the helmet repositioned such that the central focus is at the VIM.

A series of low power sonications (typically 150-300 W for ~10 seconds) produce a low temperature rise (~40-45ºC ), allowing both the neurosurgeon to assess spread of thermal field with respect to anatomy, and the neurologist to gradually assess tremor improvement.

The temperature rises are determined by planar MR thermography. Thermal maps are acquired at intervals of ~3.5 s intervals, continuing beyond the sonication to account for cooling. A sonication can be stopped by the operator (neurosurgeon), the patient (via patient alert buzzer), or by closed loop feedback within the system, including via the proprietary cavitation detection algorithm or if thermography indicates that heating exceeds the tolerable range for the given surgical stage.

Each sonication begins with a movement detection scan. If movement is detected the procedure halts until realignment is confirmed.

As sonications progress, the power delivered (and temperature at target) is gradually increased and intraoperative response monitored. When safe target engagement is confirmed, the temperature is increased to create a permanent lesion. Occasionally, based on intraoperative patient response, the focus of the transducer is slightly moved during the last sonication.

**DQA Procedure**

In the morning of each treatment day, the manufacturer recommended DQA procedure is performed. A proprietary gel phantom (Insightec, Haifa, Israel) is inserted into the transducer helmet, which is filled with water and moved into the MR bore. A series of localisation scans are performed, and test sonications begin at very low power (20 W). During each test sonication, the MR thermography plane is changed, to check the localised thermal maps are recovered in all 3 planes. The power is slowly increased as the physicist (JMcF) progresses through DQA. The electronic steering capability is checked, and finally, a higher power sonication (250 W) is delivered to check the cavitation auto-stop is working correctly. Finally, the patient ‘stop sonication’ buzzer is checked. Note that during DQA mode, the transducer output powers tested are much lower than in treatment mode, as there is no skull to absorb ultrasound.

**Treatment Setting:**

For each treatment reported in the manuscript, in the data repository at <https://osf.io/2wzqy> within the directory ‘therapeutic_ultrasound_treatment_parameters’ we provide the following as reported by the Exablate system, for each sonication contributing to the system-calculated ‘Dose Area’ (i.e. for ablative sonications):

Sonication No., Power (W), Measured Power (W), Duration (s), Actual Duration (s), Energy (J]) Measured Energy (J), Stopped (No/ {Reason}), RAS Co-ords, Actual Max Temperature (ºC), Average Max Temperature (ºC), Dose Area (mm^2^) and Treated Dose Volume (cc).

**Thermal Doses:**

We also provide, for each ablative sonication, in-house calculated maximum, average and maps of planar CEM 43 thermal doses (Sammartino et al., 2021; Sapareto & Dewey, 1984)

CEM43, the cumulative equivalent minutes at 43 ºC is a widely adopted thermal dosimetry model suitable to quantify tissue damage:

$$\mathrm{CEM}43= \int_{0}^{\tau} R^{(43-T)}dt$$

Where $R=0.5$for $T>43℃$ and $R=0.25$ for $T<43℃$.

We also provide MATLAB scripts to replicate the above data collation.

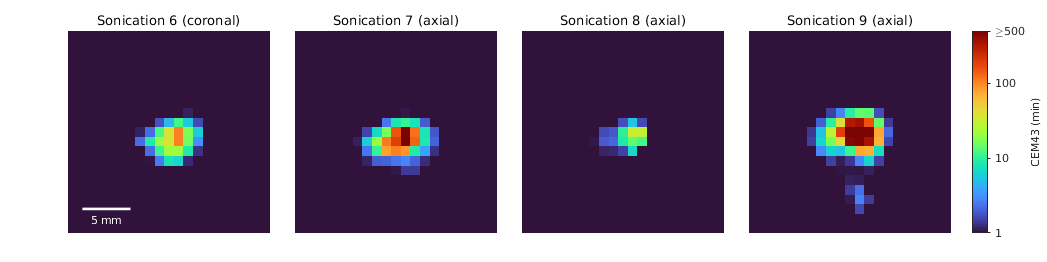

**SFig. 6: Example of CEM 43 dose maps** calculated from thermometry during ablative sonications within treatment (labelled dd_14).
